# Integrated soybean transcriptomics, metabolomics, and chemical genomics reveal the importance of the phenylpropanoid pathway and antifungal activity in resistance to the broad host range pathogen *Sclerotinia sclerotiorum*

**DOI:** 10.1101/363895

**Authors:** Ashish Ranjan, Nathaniel M. Westrick, Sachin Jain, Jeff S. Piotrowski, Manish Ranjan, Ryan Kessens, Logan Stiegman, Craig R. Grau, Damon L. Smith, Mehdi Kabbage

## Abstract

*Sclerotinia sclerotiorum*, a predominately necrotrophic fungal pathogen with a broad host range, causes a significant yield limiting disease of soybean called Sclerotinia stem rot (SSR). Resistance mechanisms against SSR are poorly understood, thus hindering the commercial deployment of SSR resistant varieties. We used a multiomic approach utilizing RNA-sequencing, Gas chromatography-mass spectrometry-based metabolomics and chemical genomics in yeast to decipher the molecular mechanisms governing resistance to *S*. *sclerotiorum* in soybean. Transcripts and metabolites of two soybean recombinant inbred lines, one resistant, and one susceptible to *S*. *sclerotiorum* were analyzed in a time course experiment. The combined results show that resistance to *S*. *sclerotiorum* in soybean is associated in part with an early accumulation of JA-Ile ((+)-7-iso-Jasmonoyl-L-isoleucine), a bioactive jasmonate, increased ability to scavenge reactive oxygen species (ROS), and importantly, a reprogramming of the phenylpropanoid pathway leading to increased antifungal activities. Indeed, we noted that phenylpropanoid pathway intermediates such as, 4-hydroxybenzoate, ferulic acid and caffeic acid were highly accumulated in the resistant line. *In vitro* assays show that these metabolites and total stem extracts from the resistant line clearly affect *S*. *sclerotiorum* growth and development. Using chemical genomics in yeast, we further show that this antifungal activity targets ergosterol biosynthesis in the fungus, by disrupting enzymes involved in lipid and sterol biosynthesis. Overall, our results are consistent with a model where resistance to *S*. *sclerotiorum* in soybean coincides with an early recognition of the pathogen, leading to the modulation of the redox capacity of the host and the production of antifungal metabolites.

**Author Summary:** Resistance to plant fungal pathogens with predominately necrotrophic lifestyles is poorly understood. In this study, we use *Sclerotinia sclerotiorum* and soybean as a model system to identify key resistance components in this crop plant. We employed a variety of omics approaches in combination with functional studies to identify plant processes associated with resistance to *S*. *sclerotiorum*. Our results suggest that resistance to this pathogen is associated in part with an earlier induction of jasmonate signaling, increased ability to scavenge reactive oxygen species, and importantly, a reprogramming of the phenylpropanoid pathway resulting in increased antifungal activities. These findings provide specific plant targets that can exploited to confer resistance to *S*. *sclerotiorum* and potentially other pathogens with similar lifestyle.

## Introduction

*Sclerotinia sclerotiorum* (Lib.) de Bary, is a plant fungal pathogen with a predominately necrotrophic lifestyle and worldwide distribution that is known to infect over 400 plant species (1). On soybean (*Glycine max* (L.) Merr.), it causes sclerotinia stem rot (SSR), a significant and challenging yield limiting disease. Data suggest that 1.6 billion kilograms of soybean are lost each year to SSR in the US alone, making it the second most damaging disease of soybean (2,3). Globally, SSR can cause yield reductions as high as 60% (4). Soybeans are an important source of plant proteins and oils globally (5–7), these products can also be significantly affected by SSR (8). Management strategies against SSR rely largely on chemical control (9), though cultural practices such as crop rotation, seeding rate and row spacing modifications have been used with limited success (3). While genetic resistance is by far more sustainable, our understanding of resistance mechanisms against this pathogen is limited, and current commercial varieties lack adequate levels of resistance to SSR.

In the past two decades, studies have increased our understanding of *S*. *sclerotiorum* pathogenic development. *S*. *sclerotiorum* is a prolific producer of cell wall degrading enzymes (CWDEs) that contribute to its pathogenic success (10). In addition to its lytic repertoire, the pathogenic success of *S*. *sclerotiorum* relies on the key virulence factor oxalic acid (OA). Mutants that are defective in OA production are weakly pathogenic (11–14). OA was proposed to contribute to pathogenesis by acidifying host tissues and sequestering calcium from host cell walls, facilitating the action of CWDEs. However, recent studies showed that *S*. *sclerotiorum* relies on OA to induce programmed cell death (PCD) in the host to its own benefit (11,15). OA induced PCD is dependent on the production of reactive oxygen species (ROS) in the plant (14,15). This points to the importance of the ROS machinery in the pathogenic success of this pathogen. Indeed, we showed that *S*. *sclerotiorum*, via OA, co-opts soybean NADPH oxidases to increase host ROS, and that the silencing of specific NADPH oxidases confers enhanced resistance to this fungus (16). In the addition to OA, other virulence factors targeting host responses are known to contribute to the pathogenicity of *S*. *sclerotiorum*. The necrosis and ethylene-inducing like proteins (NLPs) SsNEP1 and SsNEP2 induce cell death in tobacco leaves, thus contributing to disease establishment (17). The secreted chorismate mutase (SsCM1) and integrin-like protein (SsITL) were proposed to target Arabidopsis defenses by affecting salycilic acid and jasmonate signaling, respectively (18,19). The ability of *S*. *sclerotiorum* to detoxify plant chemical defenses has also been discussed (20). Unfortunately, these studies have largely focused on the fungal side of this interaction, and only provide glimpses into the plant mechanisms governing resistance to this pathogen, mainly in model plants. In soybean, Bi-parental linkage mapping led to the discovery of many quantitative trait loci (QTL) for resistance to this pathogen. Remarkably, a total of 103 QTL on 18 of the 20 soybean chromosomes have been recorded in SoyBase (21) with minimal overlap between QTL reported by different studies (9,22–26). Despite these efforts, gene level and mechanistic details of soybean resistance mechanisms against *S*. *sclerotiorum* remain unknown.

Recent advances in Next-Generation RNA sequencing (RNAseq) allow for cost-efficient and powerful examination of global differences in the transcriptional response to environmental cues. The application of RNAseq approaches in soybean-*S*. *sclerotiorum* interaction studies will, most assuredly, contribute to the development of molecular genetic resources crucial for mechanistic and translational research. Transcriptomics were used to study the interaction of *S*. *sclerotiorum* with other non-model plant hosts, including canola (27–29), pea (30), and common bean (31). While informative, these studies were solely based on gene expression comparisons, and thus may not provide a complete picture of the flow of biologic information. When coupled with other omics approaches, such as metabolomics and chemical genomics, RNAseq can provide a much greater understanding of the biology, by integrating genetic responses of a particular interaction to functional consequences. Herein, we apply multi-omics approaches to identify and validate resistance processes against *S*. *scleoriorum* in soybean lines generated in our breeding program. The identification of these processes will not only increase our understanding of the soybean-*S*. *sclerotiorum* interaction but will also facilitate the introgression of resistance into soybean varieties.

Our recent breeding efforts led to the identification of several recombinant inbred lines (RILs) highly resistant to *S*. *sclerotiorum* (9), using the soybean line W04-1002 as the main source of resistance (32). After multiple generations of greenhouse selection, we have chosen two RILs originating from the same cross; one resistant (91-145), and one susceptible (91-44) to SSR to complete this study. A comparative analysis using a combination of RNAseq, metabolomics, and chemical genomics in yeast, shows that resistance in 91-145 is associated in part with an earlier induction of jasmonate signaling, increased ability to scavenge ROS, and importantly, a reprogramming of the phenylpropanoid pathway leading to increased antifungal activities. We further discuss and provide evidence for the importance of antifungal compounds during the resistance response against this pathogen and show that this antifungal activity targets ergosterol biosynthesis in the fungus. We propose that the modulation of the identified pathways through RNAi/gene editing or overexpression approaches may be used to introgress SSR resistance in commercial soybean germplasm and possibly other host crops.

## Results

### Disease development in the resistant (91-145) and susceptible (91-44) soybean lines

To determine soybean processes involved in resistance to SSR, two recombinant inbred lines (RIL) of soybean showing a differential response to *S*. *sclerotiorum* were chosen for this study. The resistant and susceptible selections were classified based on our previous germplasm selection study (9). Both the resistant 91-145 (R) and the susceptible 91-44 (S) lines were developed utilizing the SSR resistant parental line W04-1002 (P1) and LN89-5717 (PI 5745542), an SSR-susceptible parental line having other desirable pathogen resistance traits (9). Plants were inoculated using the cut petiole inoculation method (16) and infection progression was monitored in R and S soybean lines over seven days. Initial symptoms of SSR began appearing on the main stem 48 hours post-inoculation (hpi) as light brown lesions surrounding the point of inoculation, which spread as the disease progressed. By 96 hpi, extensive tissue colonization occurred along the main stem in the S line, while small restricted lesions with a red coloration were observed at the inoculation site in the R line (Fig. 1). By day 7, it was apparent that the R line had largely restricted fungal growth on the main stem and the red coloration observed at the site of infection had become more prominent, whereas the *S*. *sclereotiorum* infection had girdled the main stem of the S line (Fig. 1). Global transcriptome analysis was conducted on *S*. *sclereotiorum* infected soybean stem tissue collected from both R and S lines at 0, 24, 48, and 96 hpi. Metabolomic analysis comparing the R and S lines was also conducted, similarly, stem tissues were collected at 0, 48 and 72 hpi.

**Figure 1.**
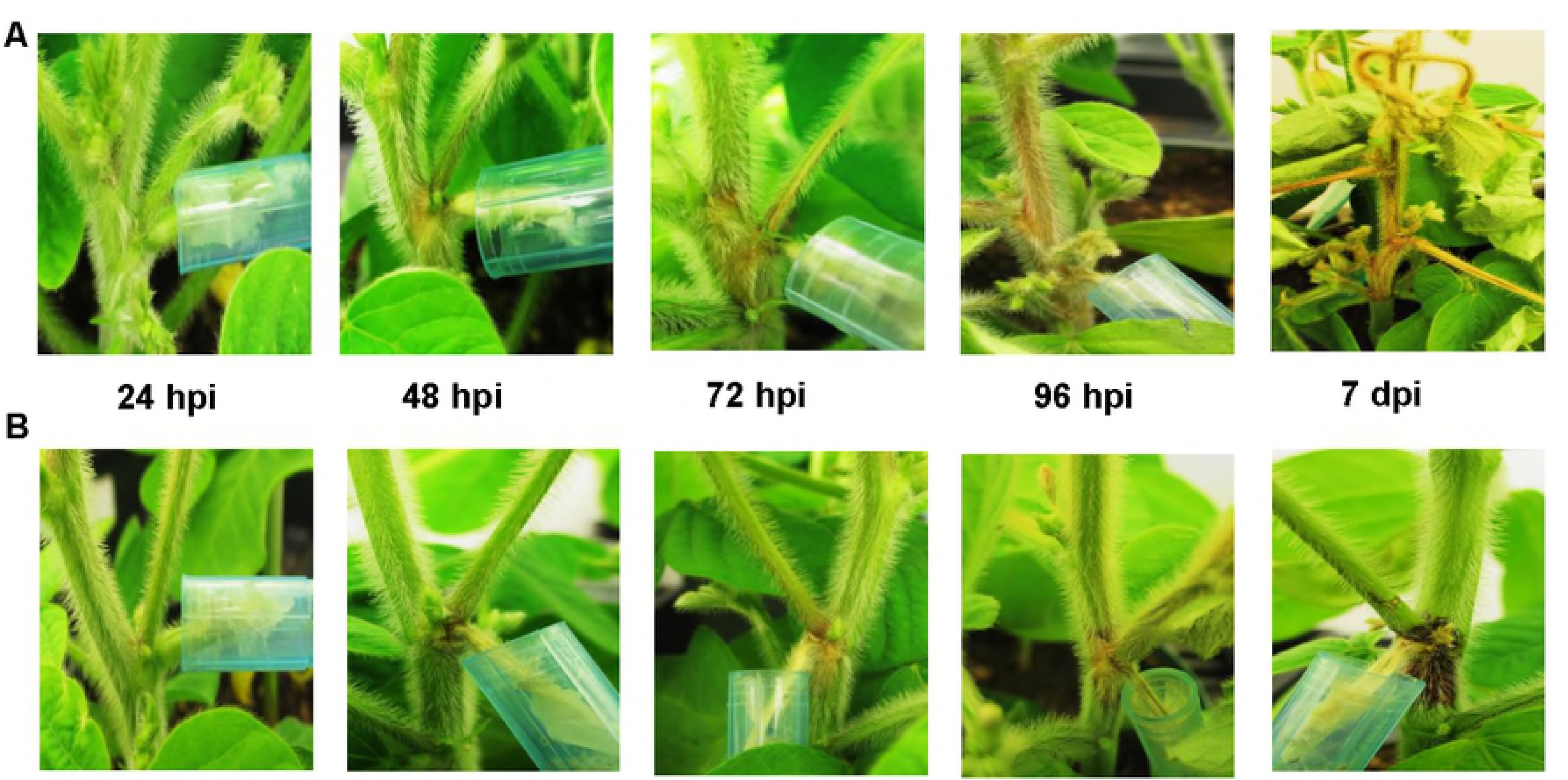
Symptom development in susceptible and resistant soybeans. Disease symptoms observed following petiole inoculation with an agar plug containing actively growing mycelia of *S*. *sclerotiorum* at 24, 48, 72, 96 hours post-inoculation (hpi) and 7 days post inoculation (dpi). (A) Susceptible (S) line. (B) Resistant (R) line. At 7 dpi in the R line, red coloration at point of inoculation (red node) is prominently visible.

### Mapping and overview of RNA sequencing data

To determine transcript levels in the R and S soybean lines over the course of infection, RNA sequencing was conducted on twenty-four stem samples, consisting of three independent biological replicates for each of the time points selected. Depending on the time point, 64.9 – 89.7 million raw reads were generated, with 95.7 - 96.6% of reads mapping to the host reference genomes of soybean and *S*. *sclereotiorum* at all timepoints. On average, 96% of the total reads mapped uniquely to the soybean reference genome in the uninfected plants of both lines. In the S line, 91.6, 91.8, and 68.9% of the reads mapped to the soybean genome at 24, 48 and 96 hpi, respectively. In the R line, 92.8, 91.9, and 88.2% of the reads mapped to the soybean genome at 24, 48 and 96 hpi, respectively (Table 1). At 96 hpi the reads mapping to the fungal genome in the S line (27%) is significantly higher than those in R line (8%), which is expected considering the extensive colonization of the soybean stem by *S*. *sclerotiorum* in the S line, particularly at the later stages of the infection process. 4.4 % to 3.2 % of reads at different time point of study did not map to any reference genome and was therefore excluded (data not shown).

**Table 1.** Summary of the sequencing metrics of the RNA-seq.

|  | **Time points**<br>**(hours)** | **Total reads** | **Mapping to**<br>***Glycine Max*** | **Mapping to**<br>***S. sclerotiorum*** |
| --- | --- | --- | --- | --- |
| **Susceptible line**<br>**(S)** | Control | 79,559,096 | 76,737,931<br>(96.5%) | 0<br>(0.0%) |
|  | 24 hpi | 81,259,978 | 74,457,988<br>(91.6%) | 3,607,921<br>(4.4%) |
|  | 48 hpi | 74,344,340 | 68,265,965<br>(91.8%) | 2,941,195<br>(4.0%) |
|  | 96 hpi | 69,266,099 | 47,765,044<br>(68.9%) | 18,712,374<br>(27.0%) |
| **Resistant line**<br>**(R)** | Control | 89,753,272 | 85,133,076<br>(96.6%) | 0<br>(0.0%) |
|  | 24 hpi | 64,923,300 | 60,261,793<br>(92.8%) | 2,286,347<br>(3.5%) |
|  | 48 hpi | 67,520,185 | 62,099,244<br>(91.9%) | 2,513,457<br>(3.8%) |
|  | 96 hpi | 72,951,207 | 64,399,890<br>(88.2%) | 5 691,551<br>(7.8%) |

### Differentially expressed genes (DEGs) of soybean during infection

A comparative analysis of differentially expressed genes (DEGs) was performed during the course of infection in both the R and S lines, as well as between these two lines at the specified time points to investigate the mechanisms underlying resistance to *S*. *sclerotiorum*. In the R line, we observed the maximum number of DEGs at 48 hours following inoculation (Table S1 and Fig. 2A). In contrast, the maximum DEGs in the S line occurred 96 hours post inoculation (Table S1 and Fig. 2B). Interestingly, at 96 hpi, the number of DEGs in the S line (14,050) were markedly higher than the R line (2,442), suggesting that the resistant response may be contributing to the reduced differential expression of host genes and the return to homeostasis at the later stages of infection. In total, 16,462 unique DEGs were identified in our lines during the course of infection (Fig. 2C). Among these, 7,319 were differentially regulated in both R and S lines, while 920 were strictly associated with the R line and 8,223 were S line specific (Fig. 2C). These results indicate that *S*. *sclerotiorum* infection causes a dramatic change in gene expression in soybean (~ 19% of total annotated genes in *Glycine max* genome), and suggests substantial differences between the resistant and susceptible soybean lines used in this study in response to *S*. *sclerotiorum* challenge.

**Figure 2.**
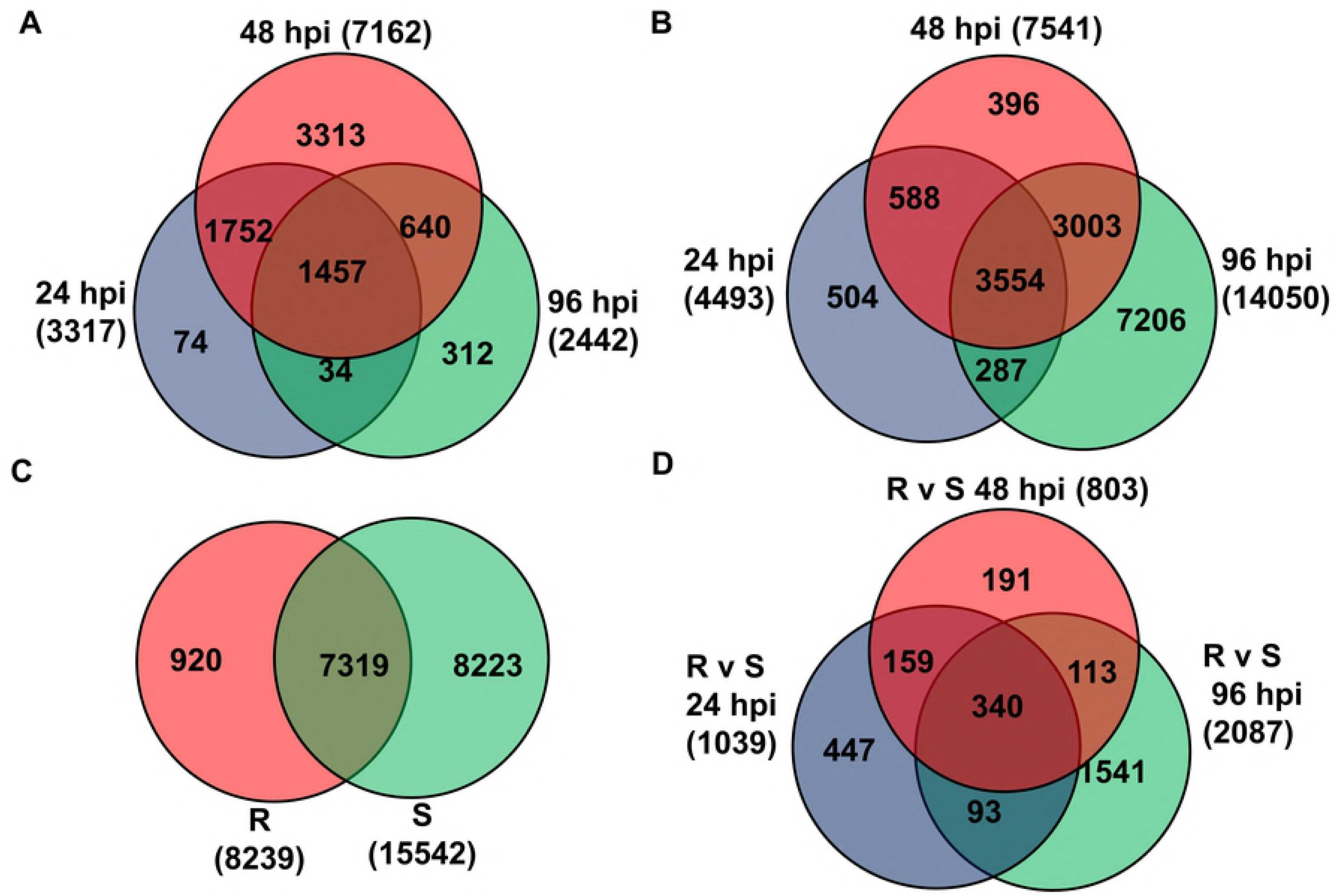
Differentially expressed genes (DEGs) identification in R and S line after *S*. *sclerotiorum* infection. Venn diagram showing (A) DEGs in R line at 24, 48 and 96 hpi compared to control (non-inoculated) sample, (B) DEGs in S line at 24, 48 and 96 hpi compared to control (non-inoculated) sample, (C) In total, 921 and 8223 DEGs were unique to the R and S line, respectively, while 7319 were identified in both the lines, (D) DEGs in R line compared to S line at 24, 48 and 96 hpi.

Pairwise comparisons of DEGs were performed between R and S soybean lines at the selected time points (Table S2 and Fig. 2D). At 24 and 48 hpi, 1,039 and 803 DEGs were identified between the two lines, respectively. However, the number of DEGs sharply increased to 2,087 at 96 hpi. This is consistent with the distinct patterns of gene expression in the S and R lines at the later stages of infection (Fig. 2 A, B). Among the identified DEGs, 447, 191, and 1541 genes were found to be time-point specific between the R and S lines at 24, 48 and 96 hpi, respectively, while 340 genes were differentially expressed at all time points following pathogen challenge (Table S2 and Fig. 2D). The putative functions of these genes were determined using Phytozome (33), National Center for Biotechnology Information (NCBI), and Soybase (21) databases (Table S2).

### Gene ontology enrichment and biological process analyses

We focus the reminder of this manuscript on direct comparisons between resistant and susceptible soybean lines, to single out potential processes associated with resistance to *S*. *sclerotiorum* in soybean. Soybean gene locus IDs identified through the DEG analysis were used to perform gene ontology (GO) enrichment analysis using the Soybase gene model data mining and analysis tool (34). A false discovery rate (FDR) value of 0.05 was used to identify significantly regulated GO biological processes, and individual GO processes were considered in this analysis if they were significantly enriched in at least one of the time points used (Table S3 and Fig. 3). The under or overrepresentation of DEGs in each GO category was determined based on the relative frequencies of the GO terms associated with the genes (34). Our data show an overrepresentation of genes in GO terms related to signal transduction (i.e. kinase signaling, phosphorylation), plant defense responses (i.e. response to chitin, fungi), phenylpropanoid pathway (i.e. chalcones, anthocyanins, flavonoids, salicylic acid), ROS scavenging, and the biosynthesis/regulation of phytohormones (i.e. jasmonic acid, salicylic acid, ethylene). DEGs underrepresented throughout the course of infection include gene related to nucleotide binding and zinc ion binding. While the involvement of categories such as ROS, defense responses, and phytohormone regulation are not surprising, our results point to complex mechanisms of gene regulation associated with resistance to *S*. *sclerotiorum*. The differential regulation of phenylpropanoid pathway genes is intriguing and will be the subject of further characterization below.

**Figure 3.**
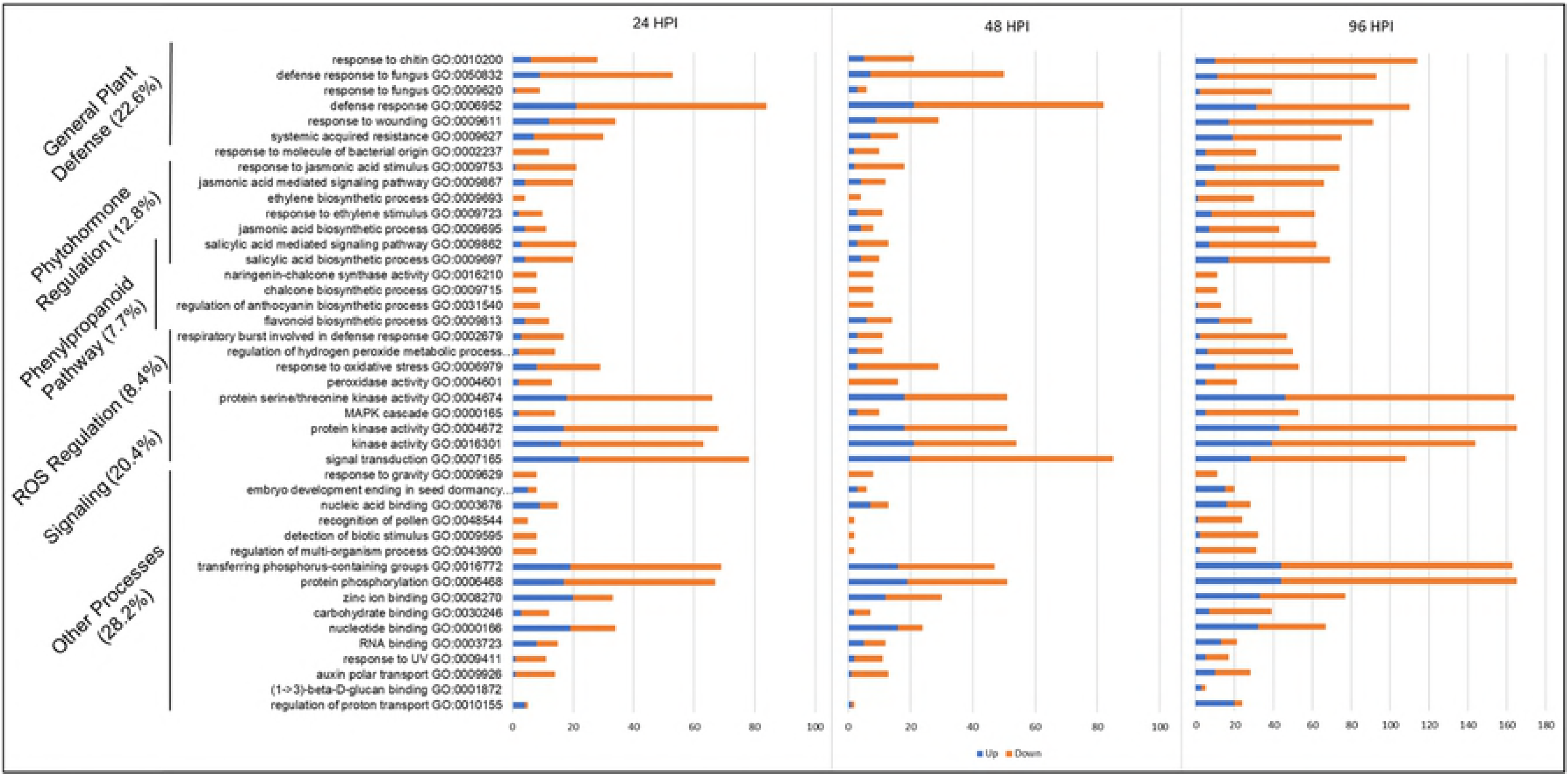
Significantly enriched Gene ontology (GO) biological processes in R line compared to S line at different time-points following infection with *S*. *Sclerotiorum*. The y-axis represents significantly enriched GO processes which were enriched (FDR <0.05) at at-least one time point of infection (24, 48 and 96 hpi). The x-axis indicates the total number of genes annotated to each GO process. Orange sections represent downregulated genes while blue sections represent upregulated genes.

### Metabolite profiling of susceptible and resistant soybean lines in response to *S*. *sclerotiorum*

Transcriptomic analysis was complemented by metabolite profiling of stem samples collected from our resistant and susceptible soybean lines following *S*. *sclerotiorum* infection. Simple changes in transcript levels might not always correlate with biological activity, but changes in metabolite flux are quantifiable outcomes that can directly explain disease phenotypes. Gas chromatography-mass spectrometry (GC-MS) analysis was performed to broadly evaluate metabolite profiles in control and infected soybean stems at 48 and 72 hpi. Three independent biological replicates were used for each time point. A total of 360 metabolites were detected, but only 164 could be identified based on available databases (Table S4). Each metabolite was characterized by its distinct retention time and mass to charge ratio (m/z). All the 164 identified metabolites found in infected soybean were also detected in non-inoculated stems and therefore are likely of plant origin. Despite this, we note that as disease progresses, contributions from *S*. *sclerotiorum* cannot be ruled out.

MetaboAnalyst 3.0 (35) was used for the analysis of the 164 identified metabolites. One-way ANOVA using Fisher’s least significant difference method (Fisher’s LSD) identified 80 metabolites, the accumulation of which was significantly (FDR <0.05) affected by *S*. *sclerotiorum* inoculation at 48 and 72 hpi in R-line compared to S-line (Table S5). These metabolites included polar compounds, such as nucleotides, amino acids, alcohols, organic acids, and carbohydrates, along with nonpolar compounds including fatty acids, and long-chain alcohols (Table S5). The multivariate analysis of identified metabolites was performed using Partial Least Squares - Discriminant Analysis (PLS-DA). The first (PC1) and second (PC2) principal components explain 52.7% of the variance. The analysis showed distinct metabolomic profiles at each time point during the course of infection, culminating with the largest segregation of metabolites between the R and S lines at 72 hpi (Fig. S1). The fold change of significantly regulated metabolites during this time course are indicated in Supplementary Table 5. Significantly regulated metabolites were assigned to distinct functional categories according to the chemical groups to which they belong (Table S5) and to specific plant pathways in which they may function (Table S6). Our data revealed several differentially regulated metabolic processes between the R and S lines (Fig. 4). However, those involved in phenylalanine metabolism are particularly interesting given the differential expression of phenylpropanoid pathway genes identified in transcriptomic analysis (Fig. 5A) and the important role of this pathway in plant defense. Indeed, the precursor of the phenylpropanoid pathway, phenylalanine, and downstream intermediate metabolites, such as benzoic acid, caffeic acid, and ferulic acid, are highly accumulated in the stem of the R line compared to the S line following *S*. *sclerotiorum* infection (Fig. 5B). Overall, the observed parallel between transcriptomics and metabolomics data with respect to the differential modulation of the phenylpropanoid pathway indicates its potential key participation in resistance to *S*. *sclerotiorum*.

**Figure 4.**
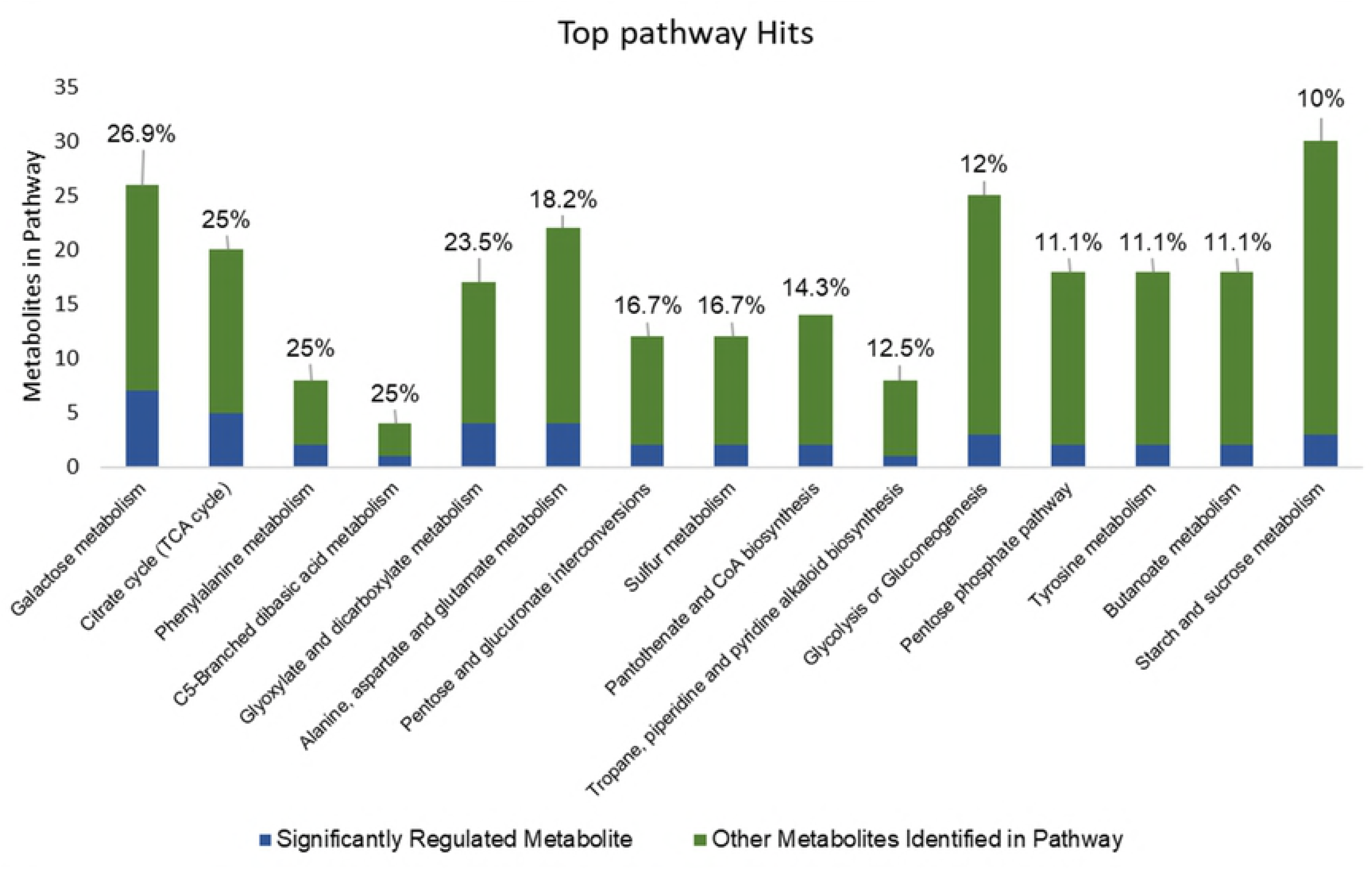
Pathway analysis of metabolites differentially detected between the R and S lines at 48 and/or 72 hpi. The y-axis represents the annotated metabolites in each pathway. Blue sections represent the differentially detected metabolites, whereas green sections represent the remaining annotated metabolites in the pathway. Percentages denote the portion of the pathway found to be differentially detected. Only pathways with at least 10% of their annotated metabolites showing significant regulation were included.

**Figure 5.**
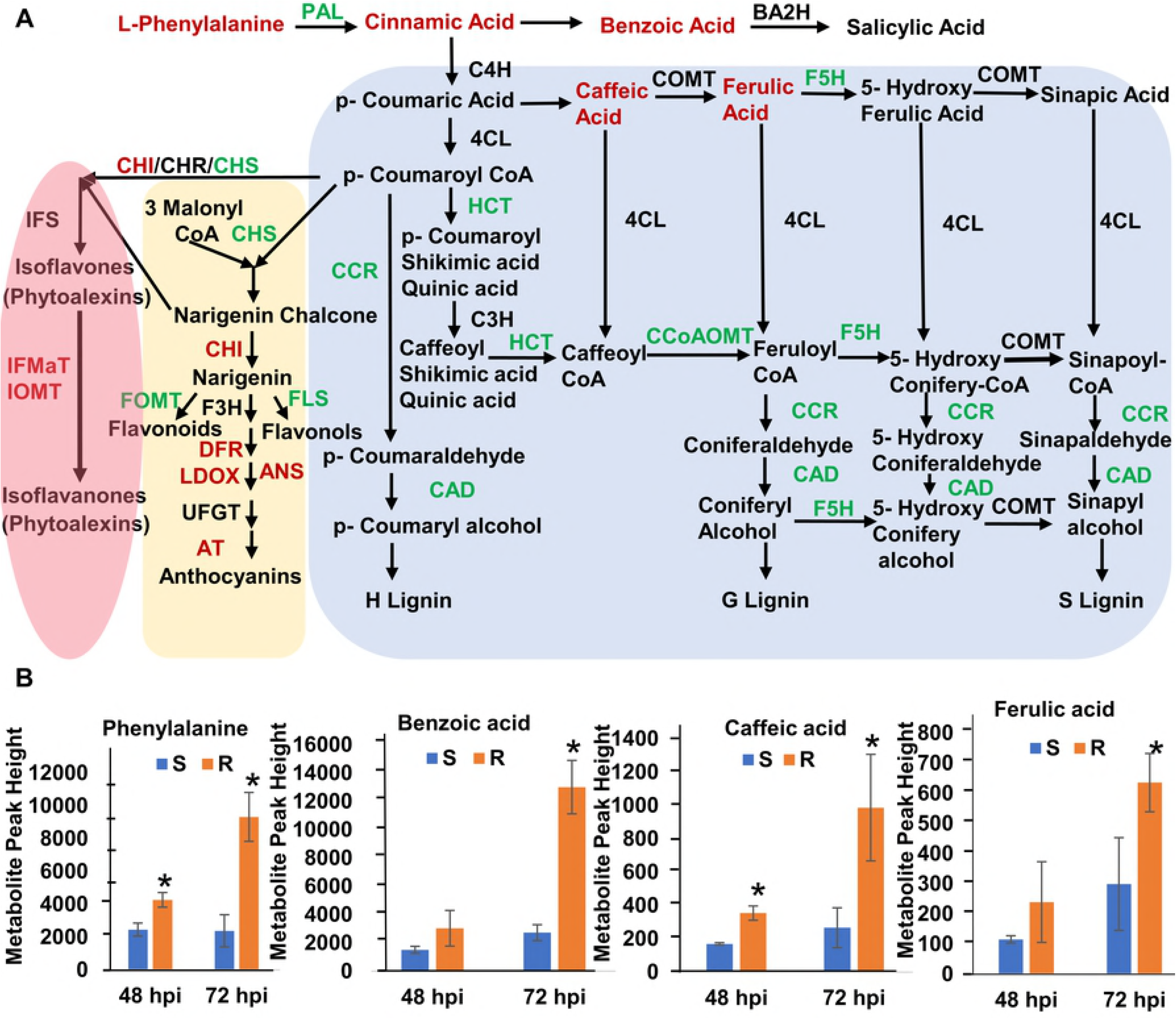
DEGs and metabolites involved in the phenylpropanoid pathway in R and S soybean lines following *S*. *sclerotiorum* infection. (A) Enzymes are indicated in uppercase letters. Gene names indicated in red or green represent significantly upregulated or downregulated genes in the R line compared to the S line, respectively. The metabolite name in red indicates significantly upregulated phenylpropanoid pathway intermediates in the R line compared to the S line. PAL (phenylalanine ammonia-lyase); BA2H (benzoic acid 2-hydroxylase); C4H (cinnamate 4-hydroxylase); COMT (caffeic O-methyltransferase); F5H (ferulic acid 5-hydroxylase); 4CL (4-coumarate: CoA ligase); C3H (p-coumarate 3 hydroxylase); HCT (N-hydroxycinnamoyl transferase); CCoAOMT (Caffeoyl-CoA O-methyltransferase); F5H (flavanone 5-hydroxylase); CCR (Cinnamoyl-CoA reductase); CAD (Cinnamoyl alcohol dehydrogenase); CHS (chalcone synthase); CHI (chalcone isomerase); CHR (chalcone reductase); FOMT (Flavonoid 4′-O-methyltransferase); FLS (Flavonol synthase); F3H (flavanone 3-hydroxylase); DHFR (dihydroflavonol-4-reductase); ANS (anthocyanidin synthase also called LDOX, leucoanthocyanidin dioxygenase); ANS (Anthocyanidin synthase); UFGT (UDP-flavonoid glucosyltransferase); AT (Anthocyanin acyltransferase); IFS (Isoflavone synthase); IFMaT (Isoflavone 7-O-glucoside-6″-O-malonyltransferase); IOMT (Isoflavone methyltransferase); this figure was adapted from Baxter and Stewart, 2013 (111) and Ferrer et al 2008 (111) (B) Estimation of the phenylpropanoid pathway intermediate metabolites (peak height) phenylalanine, benzoic acid, 3, 4 - dihydroxycinnamic acid (caffeic acid) and ferulic acid in R and S line at 48 and 72 hpi using GC – MS. Data are presented as means ± standard deviation (SD) from three independent biologically replicate with each replicate containing four pooled stem samples. ^∗^ Indicates a significant difference at p-value < 0.05 (T-Test).

Similar to transcriptomic data, the comparative metabolite profiles also implicated phytohormones in this interaction. Namely, the fatty acids linolenic acid (a precursor of jasmonic acid) and cyanoalanine (an indicator of ethylene biosynthesis) (36,37) are both significantly induced at 48 and 72 hpi, in the R line (Fig. S3). Interestingly, the most highly upregulated metabolite in our R line is mucic acid with an ~86-fold higher accumulation (Table S5). Mucic acid, also referred to as galactaric acid, can be produced by the oxidation of d-galacturonic acid, the main component of pectin.

Galactose metabolism and the TCA cycle were the pathways most affected by this analysis, and the metabolites assigned to them were primarily carbohydrates and organic acids, respectively. Interestingly, of the metabolites downregulated in R plants in comparison to S plants, 81.8% (9/11) belonged to one of these two groups (Table S5). Although these metabolites relate to multiple pathways, their downregulation in R plants may be a strategy to reduce *S*. *sclerotiorum* access to preferable carbon sources. Of the top six potentially affected pathways, three of them (Glyoxylate and dicarboxylate metabolism; Alanine, aspartate and glutamate metabolism; the TCA cycle) contain both fumaric and succinic acid. These organic acids were downregulated at 48 and 72 hpi, respectively, and demonstrate the potentially broad impacts that changes in individual metabolites can have on plant biosynthesis and metabolism.

### Reprogramming of the phenylpropanoid pathway in resistance to *S*. *sclerotiorum*

Many secondary metabolites derived from multiple branches of the phenylpropanoid pathway, including lignin, isoflavonoid-phytoalexins, and other phenolic compounds such as benzoic acid, have been proposed as important components of defense responses (38,39). In this study, we found differential expression of transcripts and metabolites related to the phenylpropanoid pathway between the R and S soybean lines. At the transcript level, we observed a downregulation of genes encoding phenylalanine ammonia-lyase (PAL), lignin biosynthetic enzymes, ferulate 5-hydroxylase (F5H), N-hydroxycinnamoyl/benzoyltransferase (HCT), Caffeoyl-CoA O-methyltransferase (CCoAOMT), cinnamoyl-CoA reductase (CCR), cinnamyl alcohol dehydrogenase (CAD), chalcone synthase (CHS), flavonol synthase (FLS), and flavonoid 4’-O-methyltransferase (FOMT) in R line compared to S line (Table 2 and Fig. 5A). Coincidentally, genes coding for enzymes involved in anthocyanin and phytoalexin biosynthesis were upregulated in the R line, these include, anthocyanidin synthase (ANS), anthocyanin acyltransferase (AT), dihydroflavonol reductase (DFR), isoflavone 7-O-glucoside-6″-O-malonyltransferase (IFMaT) isoflavone 7-O-methyltransferase (IOMT), and isoflavone reductase (IFR) (Table 2 and Fig. 5A). Transcript levels of select genes within these pathways were validated using RT-qPCR, thus confirming the RNA-Seq results (Fig. 6).

**Table 2.** A list of differentially expressed genes (DEGs) within the soybean phenylpropanoid pathway.

| Gene locus | 24 hpi<sup>a</sup> | 48 hpi<sup>a</sup> | 96 hpi<sup>a</sup> | Description |
| --- | --- | --- | --- | --- |
| Glyma.03G181700 | - | - | -1.08 | Phenylalanine ammonia-lyase |
| Glyma.03G181600 | - | -0.60 | -1.20 | Phenylalanine ammonia-lyase |
| Glyma.02G309300 | - | - | -1.28 | Phenylalanine ammonia-lyase |
| Glyma.09G186400 | - | - | -1.05 | Ferulate 5-hydroxylase |
| Glyma.04G040000 | - | - | -2.08 | N-hydroxycinnamoyl transferase |
| Glyma.01G004200 | - | - | -1.32 | Caffeoyl-CoA O-methyltransferase |
| Glyma.05G147000 | - | - | -1.54 | Caffeoyl-CoA O-methyltransferase |
| Glyma.19G007200 | - | - | -1.12 | Cinnamoyl-CoA reductase |
| Glyma.19G006900 | - | - | -1.13 | Cinnamoyl-CoA reductase |
| Glyma.09G201200 | - | - | -2.11 | Cinnamyl alcohol dehydrogenase |
| Glyma.02G130400 | - | - | -1.05 | Chalcone synthase |
| Glyma.09G075200 | - | - | -1.13 | Chalcone synthase |
| Glyma.01G091400 | - | - | -1.23 | Chalcone synthase |
| Glyma.08G110500 | -1.26 | -1.48 | -2.05 | Chalcone synthase |
| Glyma.08G110700 | -1.25 | -1.27 | -2.06 | Chalcone synthase |
| Glyma.08G109200 | -1.3 | -1.58 | -2.13 | Chalcone synthase |
| Glyma.08G110400 | -1.38 | -1.64 | -2.18 | Chalcone synthase |
| Glyma.08G110900 | -1.12 | -1.49 | -2.18 | Chalcone synthase |
| Glyma.08G110300 | -1.13 | -1.5 | -2.21 | Chalcone synthase |
| Glyma.16G219400 | - | - | -2.23 | NAD(P)H-dependent 6'-deoxychalcone synthase |
| Glyma.08G109300 | -1.17 | -1.53 | -2.24 | Chalcone synthase |
| Glyma.08G109400 | -1.3 | -1.65 | -2.42 | Chalcone synthase |
| Glyma.02G048700 | - | - | 1.49 | Chalcone-flavanone isomerase |
| Glyma.07G150900 | - | - | -2.50 | Flavonol synthase |
| Glyma.18G201900 | - | - | -3.77 | Flavonol synthase |
| Glyma.18G267800 | - | -5.27 | -9.45 | Flavonoid 4'-O-methyltransferase |
| Glyma.11G164700 | - | - | 2.99 | Dihydroflavonol reductase |
| Glyma.12G238200 | -0.88 | -0.98 | -2.07 | Dihydroflavonol reductase |
| Glyma.11G027700 | - | - | 3.27 | Anthocyanidin synthase |
| Glyma.01G214200 | - | - | 2.95 | Anthocyanidin synthase |
| Glyma.02G226000 | - | - | 1.46 | Anthocyanidin 3-O-glucosyltransferase |
| Glyma.18G271600 | 3.31 | - | - | Anthocyanin acyltransferase |
| Glyma.13G302300 | - | - | 2.49 | Anthocyanin acyltransferase |
| Glyma.13G302500 | -1.10 | - | - | Anthocyanin 5-aromatic acyltransferase |
| Glyma.14G034100 | -5.85 | -3.69 | -2.93 | Anthocyanin 5-aromatic acyltransferase |
| Glyma.19G030800 | - | - | 4.58 | Malonyl-CoA:isoflavone 7-O-glucoside-6"-O-malonyltransferase |
| Glyma.11G256500 | - | - | 4.61 | Isoflavone 7-O-methyltransferase |
| Glyma.09G094400 | - | -1.01 | - | Isoflavone-7-O-methyltransferase |
| Glyma.06G286600 | - | - | 2.69 | Isoflavone 7-O-methyltransferase |
| Glyma.13G173300 | - | - | -1.80 | Isoflavone 4'-O-methyltransferase |
| Glyma.13G173600 | - | - | -1.83 | Isoflavone 4'-O-methyltransferase |

<sup>a</sup> Fold changes (Log<sub>2</sub>FC) relative to S line.

**Figure 6.**
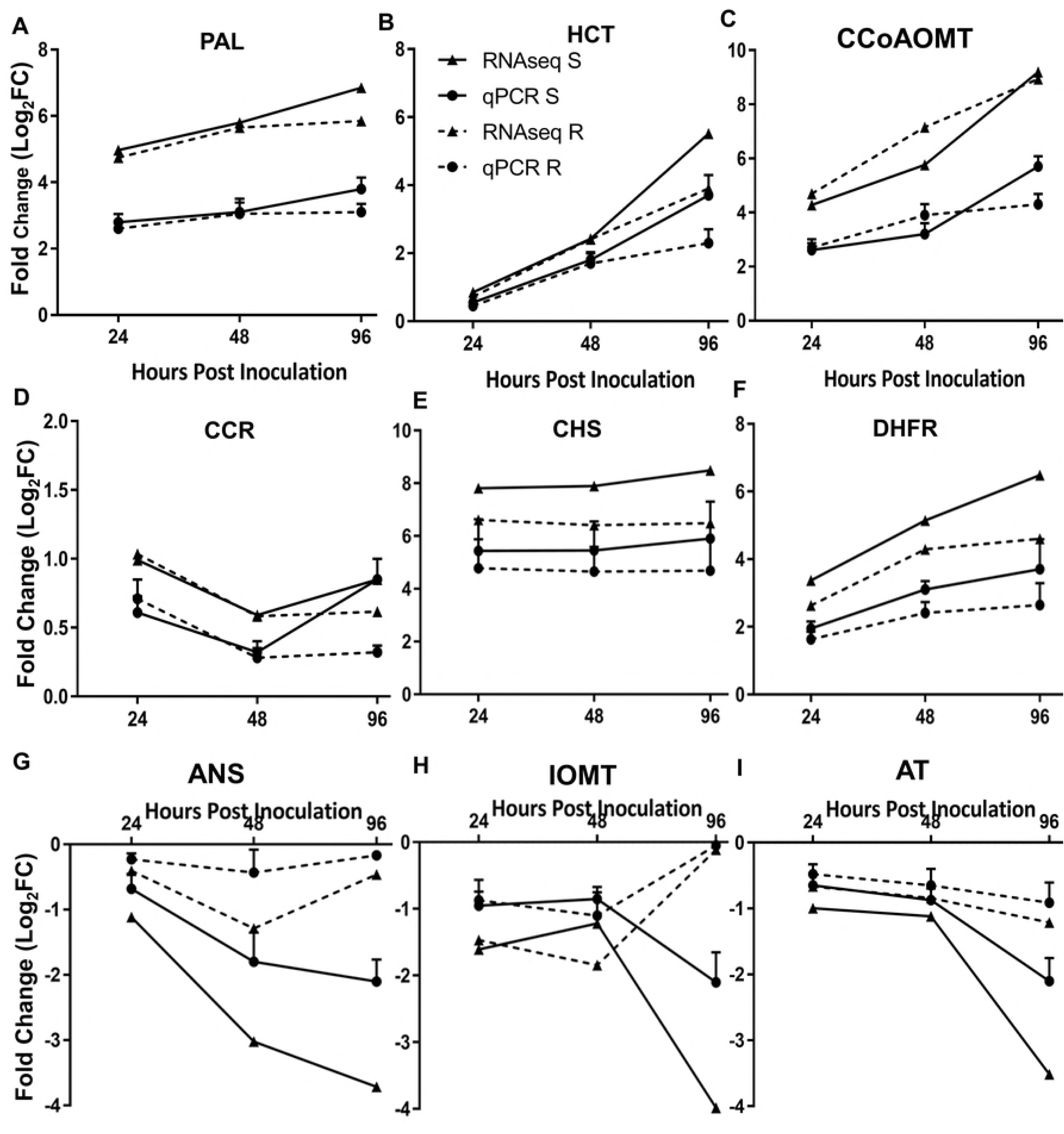
Confirmation of expression profiles of select phenylpropanoid pathway genes using qRT-PCR. (A) PAL (Phenylalanine ammonia-lyase, Glyma.02G309300); (B) HCT (N-hydroxycinnamoyl transferase, Glyma.04G040000); (C) CCoAOMT (Caffeoyl-CoA O-methyltransferase, Glyma.05G147000); (D) CCR (Cinnamoyl-CoA reductase, Glyma.19G006900); (E) CHS (Chalcone synthase, Glyma.08G110400); (F) DHFR (Dihydroflavonol reductase, Glyma.12G238200); (G) ANS (Anthocyanidin synthase, Glyma.11G027700); (H) IOMT (Isoflavone 7-O-methyltransferase, Glyma.11G256500) and (I) AT (Anthocyanin acyltransferase, Glyma.18G271600). The fold changes in expression values for qRT-PCR were calculated by comparing the expression values of genes in inoculated vs. noninfected soybean stem tissues using the 2^-ΔΔCt^ method. GmCons15 was used as endogenous control. The absolute fold changes were converted to Log_2_FC. Data are presented as means ± standard deviation (SD) from three independent experiments.

In accordance with the transcriptomics data, we observed a marked accumulation of the metabolites phenylalanine, a phenylpropanoid pathway precursor, and the lignin intermediates ferulic and caffeic acids in the R line compared to the S line (Fig. 5B). Benzoic acid, a salicylic acid precursor and known antimicrobial compound (40) also accumulated at higher levels in the Res line (Fig. 5B). Increased accumulation of ferulic and caffeic acids is likely due to the downregulation of F5H (Glyma.09G186400) and other downstream enzymes within the lignin pathway (Table 2 and Fig. 5A). We reasoned that the contribution of these ferulates to resistance may be due to their antifungal activity, and their ability to inhibit *S*. *sclerotiorum* growth was tested *in vitro*. A significant reduction of fungal growth was observed when *S*. *sclerotiorum* was grown on PDA with increasing concentrations of ferulic acid (Fig. S2). While caffeic acid did not significantly affect colony size, it clearly affected *S*. *sclerotiorum* growth patterns on PDA with the appearance of abnormal concentric ring growth patterns and premature sclerotia formation (Fig. S2).

While lignification is recognized as a disease resistance mechanism in plants, counterintuitively, our data suggest a reprogramming of the phenylpropanoid pathway away from lignin and towards the accumulation of lignin intermediates, anthocyanins, and phytoalexins in the resistance response against *S*. *sclerotiorum*. We propose that lignin intermediates, such as caffeic and ferulic acid, and possibly other compounds with antifungal activity are an important component of this response.

### ROS scavenging and antioxidant activities are associated with resistance to *S*. *sclereotiorum*

Reactive oxygen species (ROS) and ROS scavengers play active roles in redox status regulation in biotic stress (41,42). *S*. *sclerotiorum*, via oxalic acid, is known to upregulate host ROS levels to induce PCD and achieve pathogenic success (11,15,16). Our transcriptomics analysis shows a differential regulation of genes related to ROS scavenging, such as peroxidases, glutathione S-transferases (GSTs), ascorbate oxidases, and superoxide dismutase (SODs), when comparing the R and S soybean lines (Table S8 and Fig. 7). Three *GmGSTs* (Glyma.06G193400, Glyma.02G024600, Glyma.02G024800), two *GmSODs* (Glyma.12G081300, Glyma.12G178800) and five ascorbate oxidases or like proteins (Glyma.05G082700, Glyma.11G059200, Glyma.17G012300, Glyma.17G180400, Glyma.20G051900, Glyma.05G057400) were significantly upregulated in the R line compared to the S line as early as 24 hpi, suggesting a role in preventing oxidative damage imposed by *S*. *sclerotiorum* (Table S8 and Fig. 7). Peroxidases were also differentially regulated at 24 hpi, with five family members (Glyma.01G19250, Glyma.11G049600, Glyma.11G080300, Glyma.14G201700, Glyma.17G177800) significantly upregulated, however, many others were downregulated in the R line compared to S line (Table S8 and Fig. 7).

**Figure 7.**
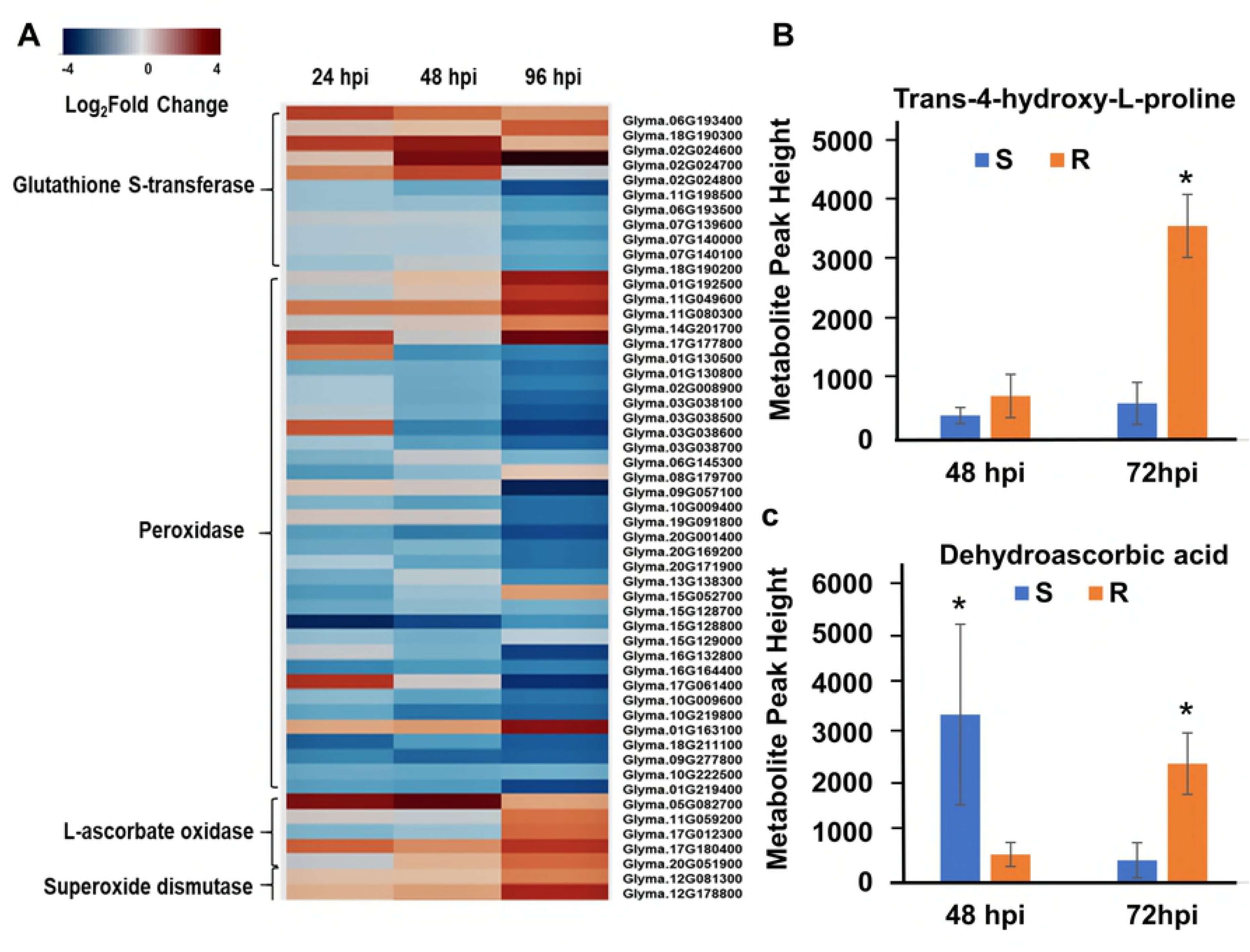
Reactive oxygen species (ROS) scavenging machinery. (A) Heat map of ROS scavenging (Glutathione S-transferase, Peroxidase, L-ascorbate oxidase, and Superoxide dismutase) and antioxidant genes (Proline-rich protein) induced in the R line compared to the S line during *S*. *sclerotiorum* infection at 24, 48 and 96 hpi. (B) Differential accumulation of ROS related metabolites trans-4-hydroxy-L-proline and dehydroascorbic acid during *S*. *sclerotiorum* infection at 48 and 72 hpi. Data are presented as means ± standard deviation (SD) from three independent experiment. ^∗^ Indicates a significantly difference at p-value < 0.05 (t-test).

We mined our metabolomics data for differentially accumulated metabolites that may serve as ROS scavengers or antioxidants. Dehydroascorbic acid (DHA), the oxidized form of ascorbate, an important antioxidant, was specifically accumulated later in the infection time course (72 hpi), but not at the early stages in the R line (Fig. 7). Similarly, the proline derivative, trans-4-hydroxyL-proline, a known osmoprotectant and antioxidant (43,44) is significantly accumulated in the R line at the later stages of the infection process (Fig. 7). Proline plays a major role as an antioxidant, owing to its ROS scavenging capacity (45,46). Overall, these results and the earlier observation of anthocyanin induction, point to a marked activation of ROS scavenging and antioxidant processes in the resistant response to *S*. *sclerotiorum*, presumably to counter to oxidative state imposed by this pathogen.

### Jasmonic acid signaling contributes to the resistant response to *S*. *sclerotiorum*

During plant-pathogen interactions, phytohormones such as salicylic acid (SA), abscisic acid (ABA), ethylene (ET), and Jasmonic acid (JA), have been shown to regulate plant immune responses (47–50). ET and JA have generally been implicated in the activation of defense responses against necrotrophs (51). Our GO enrichment analysis highlighted DEGs between the R and S lines related to JA/ET biosynthesis and responses (Table S9 and Fig. 3). Metabolic profiling also identified phytohormone-related metabolites that were differentially accumulated during the course of infection between our lines, namely, linolenic acid, a precursor of jasmonic acid, and cyanoalanine, an indicator of ethylene biosynthesis (Fig. S3). Accordingly, we conducted a targeted GC-MS analysis to more accurately estimate the dynamic changes in SA, ABA, cinnamic acid (an intermediate of SA and the phenylpropanoid pathway), and JA precursors and derivatives (12-oxophytodienoic acid (OPDA), and (+)-7-iso-jasmonoyl-L-isoleucine (JA-Ile) in a time course experiment comparing the R and S soybean lines. Distinct patterns of phytohormone accumulation were identified during the course of infection (Fig. 8). The bioactive form of JA, Jasmonoyl-L-isoleucine (JA-Ile) but not JA, was significantly induced in the R line compared to the S line between 6 and 24 hpi, before decreasing at the later stages of our time course (Fig. 8). Interestingly, JA and JA-Ile levels drastically increased in the S line, albeit at a much later stage of infection (48 – 72 hpi). Thus, this late dramatic surge in JA and JA-Ile in the S line may be perceived as a delayed response to fungal colonization, perhaps after the establishment of infection (Fig. 7). In accordance, OPDA, a precursor of JA, accumulated at significantly higher levels in the S line at 48 – 72 hpi (Fig. 8). This pattern also explains the significant induction of JA biosynthetic transcripts at the later stages of infection observed in the S line (Table S9). Expectedly, SA accumulated to higher levels in the S line throughout the time course (Fig. 8). JA and SA responses are known to be antagonistic in many plant-pathogen interactions (52). *In toto*, phytohormone estimation in the S and R lines following *S*. *sclerotiorum* challenge suggests that resistance to this pathogen in soybean coincides with an early induction of JA signaling. The timing of this induction appears to be critical to the outcome of this interaction.

**Figure 8.**
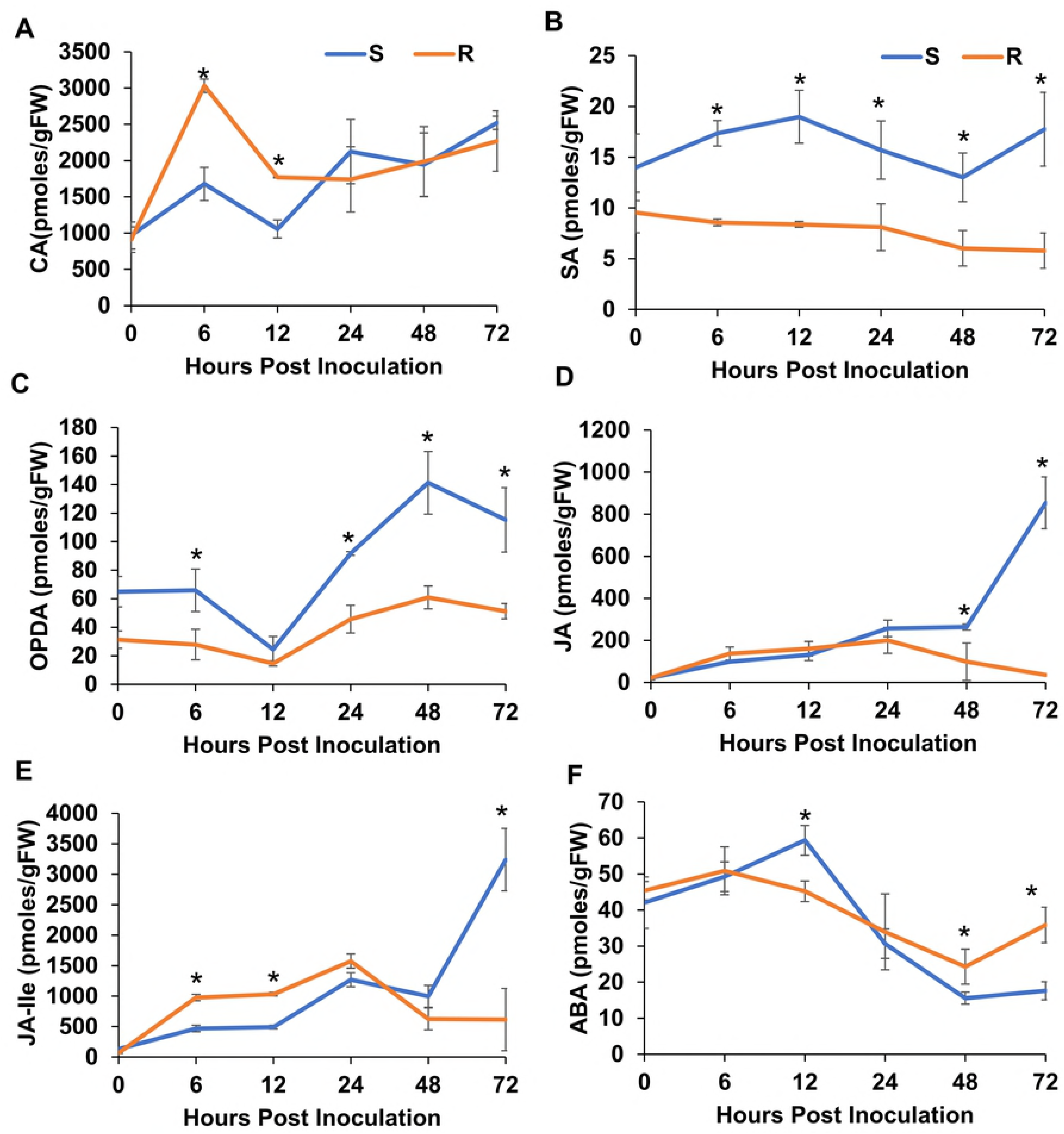
Estimation of phytohormones and a phenylpropanoid pathway precursor (Cinnamic acid, CA) in R and S lines following *S*. *sclerotiorum* infection. (A) Cinnamic Acid (CA), (B) Salicylic Acid (SA), (C) Abscisic Acid (ABA), (D) Jasmonic Acid (JA), (E) JA precursor 12-oxophytodienoic acid (OPDA) and (F) bioactive JA derivative (+)-7-iso-jasmonoyl-L-isoleucine (JA-Ile). The bars represent the standard deviation (n = 3). ^∗^ Indicates a significant difference between the R and S lines at p-value < 0.05 (One-way ANOVA).

### Antifungal activity and mode of action of stem extract from the resistant soybean line

Our earlier observation that soybean stems from the R line accumulated metabolites with antifungal activity, such as ferulic and caffeic acids, following *S*. *sclerotiorum* challenge was intriguing. This resistant response was also associated with the development of a prominent red coloration at the site of infection (Fig. 1). Considering the metabolomics data, we reasoned that total stem extract from the resistant line should also exhibit antifungal activity against *S*. *sclerotiorum*. To test this hypothesis, we prepared an ethanol extract of stem sections harvested from the R line,10 days post inoculation, that we termed the red stem extract. Equal amount of green stem extracts harvested from non-inoculated plants served as control. The final extracts were diluted in DMSO following ethanol evaporation. *S*. *sclerotiorum* growth was assayed on potato dextrose broth (PDB) amended with the red stem extract, green stem extract, or DMSO control for 48 hours. Fungal biomass as determined by mycelial fresh weight was markedly reduced (12-14 fold) in PDB cultures containing the red stem extract compared to PDB amended with the green stem extract or DMSO (Fig. S4 B, C). These results confirm that the resistant response associated with our R line clearly involves the accumulation of antifungal compounds that inhibit *S*. *sclerotiorum* growth.

We next examined the mechanism by which the red stem extract inhibits fungal growth by performing chemical genomic profiling in yeast. Chemical genomic profiling is built upon barcoded strains of *Saccharomyces cerevisiae* mutants representing approximately 4000 gene deletions. The use of unique barcodes for each mutant makes this approach compatible with high-throughput screening of drugs using next-generation sequencing (53). Chemical genomic profiling revealed that deletion mutants of genes involved in phospholipid and sterol biosynthesis were significantly sensitive to the red stem extract (Fig. 9A). Mutants of *ERG6*, which encodes a protein in the ergosterol biosynthetic pathway, had the greatest sensitivity to the extract, and this was a highly significant response (p<1e-7). *ERG2*, which is also involved in ergosterol biosynthesis, was also significantly sensitive (p<0.01). Mutants of *ARO7*, which encode a gene involved in amino acid biosynthesis was significantly sensitive. While *ARO7* is not known to be directly involved in lipid/sterol biosynthesis, it has many genetic interactions with lipid related genes (54). *CHO2* and *OPI3* mutants were also significantly sensitive. Cho2p and Opi3p are both involved in phosphatidylcholine biosynthesis. A deletion mutant of *PAH1* was the most significantly resistant mutant. Pah1p is a phosphatase that regulates phospholipid synthesis. Deletion mutants of *PAH1* have increased phospholipid, fatty acid and ergosterol ester content (55). Further, in two out of three replicates, the chemical genomic profile of the red stem extract had significant correlation (p<0.05) with the profile of fenpropimorph (56), an ergosterol biosynthesis inhibitor that targets *ERG2* and *ERG24* in yeast. Taken together, these data suggest that the red stem extract may exert toxicity by either disrupting enzymes involved in lipid/sterol biosynthesis, or alternatively physically binding membrane lipids and causing cell leakage.

**Figure 9.**
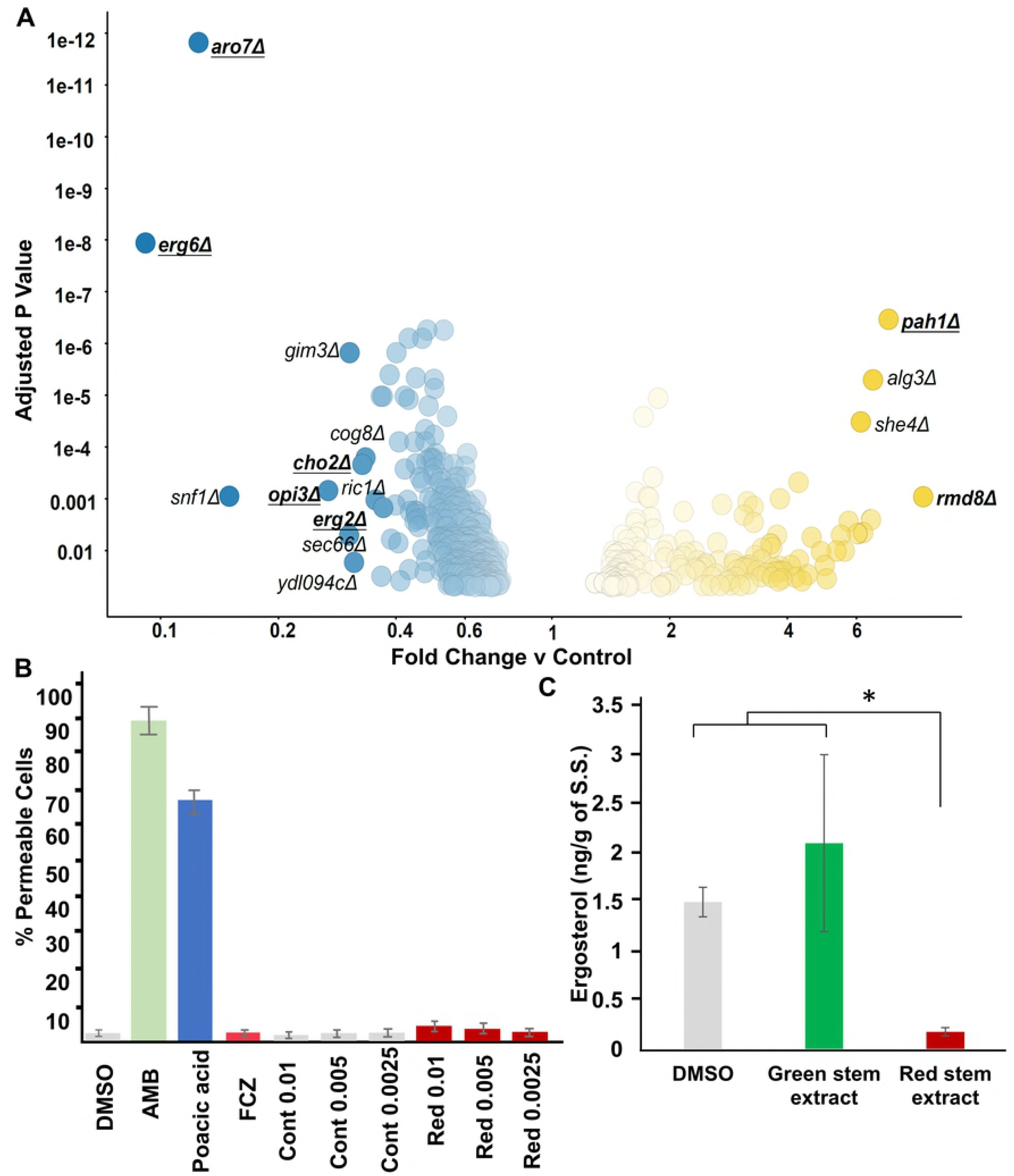
Chemical genomic profiling of the red stem extract in yeast mutants and ergosterol biosynthesis. (A) Plot showing growth differences between yeast deletion mutants exposed to the red stem extract. Dots represent mutants which were significantly sensitive (blue) or resistant (yellow) to the fungicidal activity of the extract. Underlined mutants are implicated in phospholipid and sterol biosynthesis (B) cell leakage assay to test the permeability of *S*. *sclerotiorum* treated with DMSO, fluconazole (FCZ), amphotericin B (AMB), poacic acid, and different concentrations of green (Cont) and red (Red) stem extract, (C) ergosterol estimation from *S*. *sclerotiorum* treated with DMSO, green stem extract and red stem extract. Vertical bars show standard deviation of means of three replicates. ^∗^ Indicates a significant difference at p-value < 0.05 (t-test).

**Figure 10.**
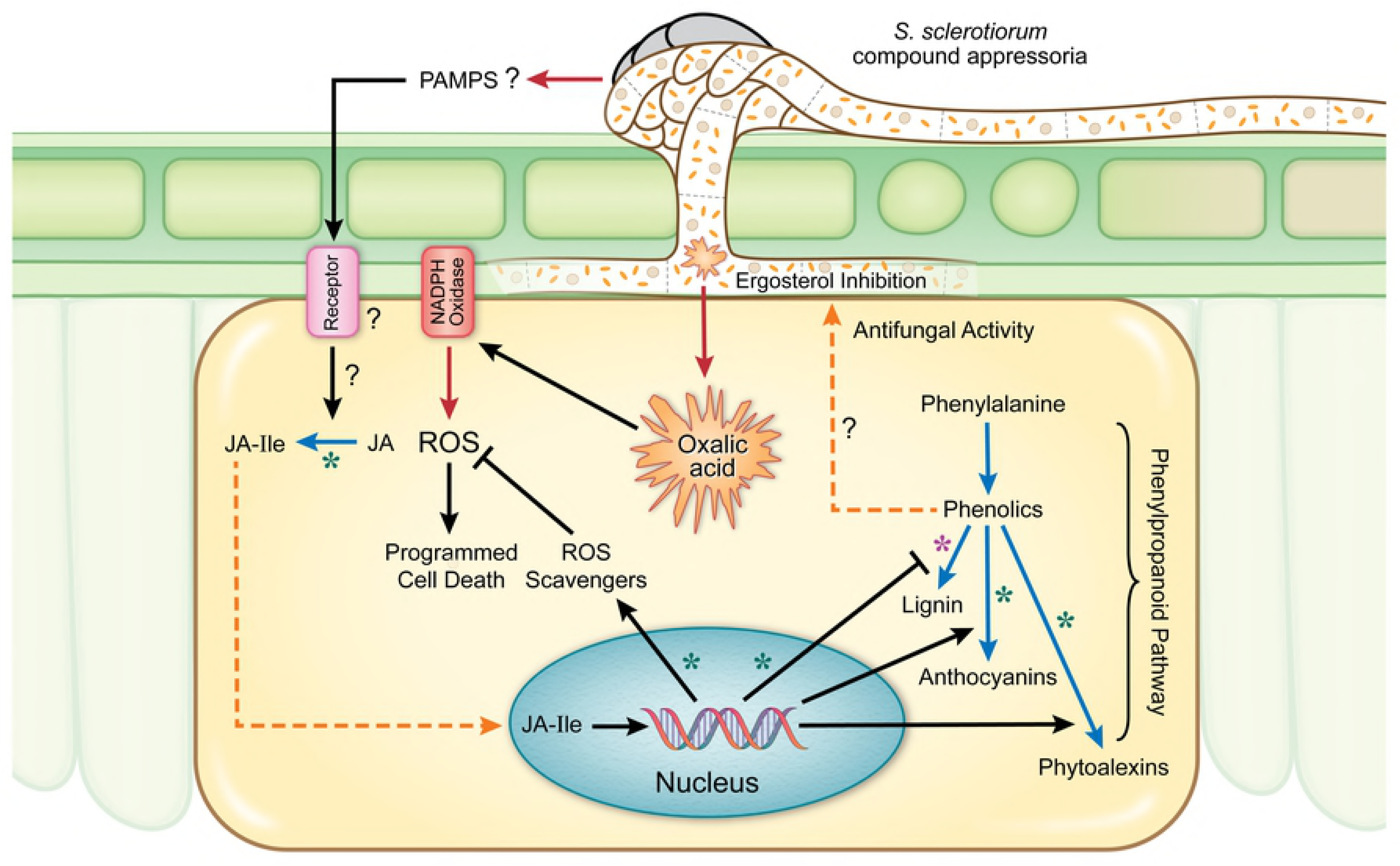
Cellular model summarizing *S*. *sclerotiorum* resistance mechanisms in soybean. Black line = Induction/suppression of a process, Red line = Secretion/release, Blue line = Bioconversion, Dashed orange line = Translocation of a metabolite, purple asterisk (^∗^) = Downregulation in resistant response, Green asterisk (^∗^) = Upregulation in resistant response, JA = Jasmonic Acid, JA-Ile = Jasmonic Acid-Isoleucine.

### Red stem extract inhibits ergosterol biosynthesis in *S*. *sclerotiorum*

To test if the red stem extract causes rapid cell lysis like the antifungal compounds amphotericin B (binds ergosterol) and poacic acid (binds glucan), we performed a cell leakage assay. The red stem extract did not cause significant cell permeability (Fig. 9B). The antifungal drug fluconazole inhibits ergosterol biosynthesis but doesn’t cause rapid cell permeability as it targets the enzyme Ergllp rather than a membrane or cell wall structure. This result supports the hypothesis that compounds within the red stem extract target ergosterol biosynthetic enzymes. We thus tested if treatment of *S*. *sclerotiorum* with the red stem extract alters ergosterol production. *S*. *sclerotiorum* was grown in PDB amended with red stem extract, green stem extract, or DMSO, and ergosterol level was calculated by first calculating the total ergosterol-plus-24(28) DHE (dehydroergosterol) content and then subtracting from the total the amount of absorption due to 24(28) DHE only (57). Ergosterol content of *S*. *sclerotiorum* mycelia grown in red stem extract was significantly reduced compared to control treatments (Fig. 9C). The results confirm that the red stem extract inhibits the growth of *S*. *sclerotiorum* by targeting ergosterol biosynthesis in the fungus.

## Discussion

Resistance to fungal pathogens with a predominately necrotrophic lifestyle, such as *Sclerotinia sclerotiorum*, is not well understood due to the likely complex network of responses to these pathogens or their determinants. Uncovering key components of these defense responses is essential for the deployment of disease resistant crops. Omics approaches offer a unique opportunity to identify global cellular networks in plants in response to these pathogens. *S*. *sclerotiorum* is a broad host pathogen that infects over 400 species, mostly dicotyledonous plants. Its pathogenic success most assuredly relies on a broadly effective toolkit that allows it to infect multiple hosts, however, host-specific dialogue between a given host and the pathogen may also be of importance. In this study, we specifically examine soybean resistance mechanisms against *S*. *sclerotiorum* by comparing opposing outcomes in two soybean lines in response to this pathogen using a combination of transcriptomics, metabolomics, and chemical genomics. Several lines of evidence are consistent with the following conclusions: (i) *S*. *sclerotiorum* challenge induces drastic changes in gene expression and metabolite production in soybean; (ii) Resistance in soybean is associated with an early recognition of the pathogen and a rapid induction of JA signaling; (iii) The redox buffering capacity of the host is essential to counter the oxidative state imposed by *S*. *sclerotiorum*; (iv) A reprogramming of the phenylpropanoid pathway and upregulation of antifungal metabolites are observed during the resistant response to *S*. *sclerotiorum*; (v) The antifungal activity associated with resistance targets ergosterol biosynthesis in the pathogen. Overall, comprehensive genetic, biochemical, and transcriptomic analyses allowed us to uncover a novel resistance mechanism connecting the upregulation of antifungal activity to a successful defense response against *S*. *sclerotiorum* and highlights the importance of early recognition and redox regulation in resistance to this pathogen.

The importance of ROS in plant immunity and other plant processes, including abiotic stress responses, growth and development, is well documented (58). In plant immunity, ROS can function <u>not only</u> as antimicrobials <u>and</u> in plant cell wall reinforcement, but also as signaling molecules activating additional defense responses (59). The implication of host ROS in plant defenses, including the hypersensitive response (HR) and pathogen-associated molecular pattern (PAMP)-triggered immunity following pathogen recognition is well documented (60,61). However, ROS are also produced during compatible interactions, thus facilitating host colonization of certain fungal pathogens (11,14,16,62). *S*. *sclerotiorum* is known to induce apoptotic-like PCD via oxalic acid, a process that requires ROS upregulation in the host (15). Indeed, we have recently shown that this pathogen hijacks soybean NADPH oxidases to increase ROS levels leading to tissue death and the establishment of disease, and that the silencing of specific host NADPH oxidases confers enhanced resistance to this pathogen (16). This study confirms the importance of ROS in this pathosystem and suggests ROS scavenging and antioxidant activity as viable resistance mechanisms in soybean. Indeed, we noted the significant upregulation of genes related to ROS scavenging such as peroxidases, glutathione S-transferases, ascorbate oxidases, and superoxide dismutase, in association with the successful defense response against *S*. *sclerotiorum*. Similar trends were reported in other host and non-host plant species in association with *S. sclerotiorum*, with strong upregulation of genes encoding ROS scavenging enzymes and higher antioxidant enzymatic activities coinciding with resistance to this pathogen (63–66). We also noted the accumulation of antioxidant metabolites dehydroascorbic acid (DHA) and trans-4-hydroxy-L-proline. DHA is converted into ascorbic acid (AA), which is known for its redox buffering capacity and ROS detoxification (67,68). The utilization of ascorbic acid as an antioxidant in cells causes its oxidation back to dehydroascorbic acid (69). Thus, the low levels of DHA in our resistant line at the onset of infection can be explained by higher ascorbic acid levels at this stage. <u>O</u>nce oxidized, AA is converted to DHA, which accumulated at the later stages of our time course. Similarly, the proline derivative, trans-4-hydroxy-L-proline, was markedly increased in our resistance response, and is a known osmoprotectant and antioxidant (43,44). The accumulation of proline in plants has been implicated in stress tolerance by maintaining osmotic balance, stabilizing membranes, and modulating ROS levels (43). Overall, our accumulating evidence suggests that the antioxidant capacity of soybean plays a critical role in its ability to resist *S*. *sclerotiorum*, a pathogen that induces ROS and cell death in the host to achieve pathogenic success.

An important question is what causes this stark difference in ROS buffering capacity between the R and S soybean lines despite common genetic components? We propose that the early recognition of the pathogen in the R line leads to a timely response that includes the activation of the antioxidant machinery within the host. This is corroborated by our phytohormone analysis that shows a rapid induction of the bioactive form of jasmonic acid (JA), Jasmonoyl-L-isoleucine (JA-Ile) as early as 6 hpi in the resistance response, JA-Ile levels decreased at the later stages of infection, presumably once the infection was under control. In contrast, in the susceptible response, JA and JA-Ile levels remained low early, but drastically increased at the later stages of our time course, which can be conceived as a delayed and unsuccessful response to fend off an already established infection. A large body of work on plant defenses described the implication of specific hormone pathways depending on pathogen lifestyle (47–50,52). Plant defenses involving JA typically inhibit fungal necrotrophs (52), and mutants specifically impaired in JA-Ile accumulation show enhanced susceptibility to such pathogens (70–72). Our results are consistent with the model where upon *S*. *sclerotiorum* challenge, JA is rapidly biosynthesized from linolenic acid and subsequently catalyzed to JA-Ile, the active form of JA. Our metabolomics analysis also showed that linolenic acid is hyper accumulated early in the resistance response. JA-Ile is known to act upon the F-box protein COI1 (Coronatine-insensitive protein 1) leading to the targeting of JAZ (Jasmonate-Zim Domain) proteins for degradation, thus liberating JAZ repressed transcription factors involved in defense (72,73). Thus, while JA signaling appears to be activated in both resistant and susceptible responses, the timing of this induction is key to the resistance outcome in *S*. *sclerotiorum*-soybean interaction. The early induction likely leads the timely activation of defense components culminating in the arrest of fungal growth and colonization. The reciprocal antagonism between JA and SA signaling pathways is often discussed, and SA signaling is expected to have a negative effect on resistance to pathogens with a predominately necrotrophic lifestyle (52). Our data indicates a higher and sustained levels of SA in the susceptible line throughout our time course, thus in line with the notion that these signaling pathways are antagonistic. SA is also associated with elevated ROS levels and cell death induction, to the benefit of necrotrophic pathogens (74). However, the exact role of SA in this interaction will require further investigation knowing that the crosstalk between JA and SA signaling is complex, and synergistic interactions have also been reported (50).

JA signaling have been linked to the alteration of plant secondary metabolites, including alkaloids, terpenoids, flavonoids, phenolic compounds, and phytoalexins (75–79). Secondary metabolites derived from multiple branches of the phenylpropanoid pathway, including lignins, isoflavonoid-phytoalexins, and other phenolic compounds have also been proposed as important components of defense responses (38,39). The integration of our transcriptomic and metabolic data revealed that the phenylpropanoid pathway is differentially regulated in our soybean lines in response to *S*. *sclerotiorum* challenge. Specifically, our data suggest a reprogramming of the phenylpropanoid pathway with its flux diverted from lignin to lignin intermediates, anthocyanins, and phytoalexins in the resistance response against *S*. *sclerotiorum*. While these results will require further confirmation by the absolute quantification of phenylpropanoid pathway components, the accumulation of upstream metabolites such as ferulic acid, caffeic acid, and benzoic acid in the resistance line support a reduced flow towards lignins. The observed accumulation of ferulic acid is also consistent with decreased lignification considering its role as a nucleation site for lignin polymerization (80). These results may seem counterintuitive considering that lignin biosynthesis is associated with cell wall fortification as a mechanism of disease resistance (81). However, in support of our results, a negative correlation between soybean stem lignin content and resistance to *S*. *sclerotiorum* has previously been reported (32). Flux changes within the phenylpropanoid pathway in maize have been discussed in response to the biotrophic fungal pathogen *Ustilago maydis* (82,83). Interestingly, *U*. *maydis*, via the secreted effector Tin2, diverts the flow of the phenylpropanoid pathway away from lignins by increasing anthocyanin biosynthesis to facilitate infection. In the absence of this effector, lignin biosynthesis is enhanced presumably to limit fungal colonization (83). In contrast to our results, resistance to *U*. *maydis* appears to be associated with the activation of the lignin branch of the phenylpropanoid pathway. However, these observations are in line with the limited lytic repertoire and the biotrophic lifestyle of *U*. *maydis*. Against necrotrophic fungal pathogens, anthocyanin may act as antioxidants by scavenging ROS and limiting the induction of cell death required by these pathogens. Indeed, the ROS scavenging capacity of anthocyanins has been proposed to provide protection against necrotrophic pathogens such as *Botrytis cinerea* (84) and *Erwinia carotovora* (85). We propose that this mechanism may also confer resistance against *S*. *sclerotiorum* in soybean.

The reported antimicrobial activities of ferulic (86), caffeic (87), and their significant accumulation in the resistance response against *S*. *sclerotiorum* prompted us to consider antifungal activity as a component of the soybean defense response against this pathogen. Indeed, total red stem extracts from our resistant line following *S*. *sclerotiorum* challenge clearly inhibited fungal growth *in vitro*. This antifungal response was seemingly absent from healthy soybean plants, activated only in response to *S*. *sclerotiorum*. Furthermore, we were able to show using chemical genomics in yeast that factors within this antifungal activity target ergosterol biosynthesis in the fungus. Ergosterol, a lipid found in the cellular membranes of fungi, is important to the regulation of membrane fluidity and permeability. It is therefore conceivable that plants have evolved means to target this key component of fungal membranes. For example, saponins, which affect membrane integrity by tergeting 3β-hydroxyl sterols, were shown to be required for resistance against *Gaeumannomyces graminis* var. *tritici* and several *Fusarium spp* (88–90). Plant antimicrobial peptides and coumarin, a product of the phenylpropanoid pathway, were also proposed to target fungal ergosterol (91,92). The specific metabolites responsible for these antifungal activities in our resistant line are currently unknown. While we provided evidence that both ferulic and caffeic acids affected *S*. *sclerotiorum* growth, their chemical genomic profile did not match that of the red stem extract. Thus, we propose that other unknown compounds target ergosterol biosynthesis and contribute to resistance to this pathogen. The identification of these compounds through high-resolution mass spectrometry may lead to the discovery of novel bioactive metabolites and help devise specific strategies to introgress resistance to fungal pathogens in crop plants.

## Methods

### Plant material and pathogen inoculation

Two recombinants inbred lines of soybean (*Glycine max*), 91-145 and 91-144, were used in this study. Both the resistant 91-145 (R) and the susceptible 91-44 (S) lines were developed utilizing W04-1002 (P1), a SSR resistant parental line, and LN89-5717 (PI 5745542), a SSR-susceptible parental line demonstrating other desirable pathogen resistance traits (9). Soybean seedlings and plants were maintained in the greenhouse or growth chamber at 24 ± 2°C with 16-h light/8-h dark photoperiod cycle. Plants were supplemented fertilizer (Miracle-Gro) every two weeks.

SSR infection was performed using a wild type strain of *S*. *sclerotiorum* (1980) grown at room temperature on potato dextrose agar (PDA) as described by Godoy et al. 1990. Four-week-old soybean plants were infected with *S*. *sclerotiorum* by petiole inoculation, using an agar plug of actively growing fungal hyphae. Plant tissue was sampled by cutting horizontally above and below (1.5 cm) the node of the inoculated petiole with a clean straight-edge razor (16). Tissue samples were then immediately frozen in liquid nitrogen prior to RNA extraction and metabolomic analysis. Samples from non-inoculated stem tissues were also collected as a control. The experimental design was completely randomized and consisted of three biological replicates for each of the treatments. Each biological replicate consisted of stem segments (~3 cm, first internode) from 2 different plants.

### RNA Extraction and library preparation

Total RNA was extracted from soybean stem tissues using a modified Trizol protocol (Invitrogen Corp., Carlsbad, CA, USA). Briefly, collected tissue from each sample was finely ground in liquid nitrogen. For each 100 mg of tissue, 1 ml of chilled Trizol was added. Samples were centrifuged at 12k rpm for 5 min at 4°C. Supernatant was discarded and 200 μ1 of chilled chloroform was added, and vortexed at high speed for 15 sec. Samples were centrifuged again at 12k rpm for 15 min at 4°C. The aqueous phase was mixed with 0.8x isopropanol and left at room temperature for 10 minutes, followed by centrifugation at 12k rpm at 4°C. Supernatant was discarded, and the pellet was washed with 75% ethanol. Pellet was air dried for 10 minutes at room temperature and resuspended in 20μl of nuclease free water followed by incubation at 55°C for 10 minutes. Samples were cleaned using the RNeasy Plant Mini Kit (Qiagen, Hilden, Germany). RNA concentration and purity were determined by Nanodrop (Thermo Fisher Scientific, Wilmington, DE) and sample quality was assessed using an Agilent Bioanalyzer 2100 and an RNA 6000 Nano Kit (Agilent Technologies, Santa Clara, CA). The RNAseq experiment included three biological replicates per treatment.

Library preparation was performed at the University of Wisconsin – Madison Biotechnology Centre (Madison, WI, USA). Individually indexed libraries were prepared using the TruSeq RNA Sample Preparation v2 kit according to the manufacturer’s instructions (Illumina, San Diego CA, USA). Library concentrations were quantified with the Qubit HS DNA kit (Thermo Fisher Scientific, Wilmington, DE). The size and quality of the libraries were evaluated with an Agilent Bioanalyzer 2100 and an Agilent DNA 1000 kit (Agilent Technologies, Santa Clara, CA) and the libraries were sequenced using Illumina HiSeq2500 (1X100bp) (Illumina, San Diego CA, USA).

### Quality check and sequence analysis

Illumina raw read data quality was verified with fast QC. The soybean and *S*. *sclereotiorum* genome sequences were acquired from Phytozome v12.1 (https://phytozome.jgi.doe.gov/pz/portal.html#!bulk?org=Org_Gmax) and the Broad institute (https://www.broadinstitute.org/fungal-genome-initiative/sclerotinia-sclerotiorum-genome-project), respectively (10,93). Raw sequence reads were mapped to both genomes using the Subjunc aligner from Subread (94). Alignments were compared to the gene annotation GFF files for both organisms (Soybean: Gmax_275_Wm82.a2.v1.gene.gff3 (93), *S*. *sclereotiorum*: sderotinia_sderotiorum_2_transcripts.gtf (10)) and raw counts for each gene were generated using the feature Counts tool from subread. The raw counts data were normalized using voom from the R Limma package, then used to generate differential expression (log_2_FC) values (95,96). DEGs were generated from the comparison of inoculated soybeans of both lines at different time points to their respective uninoculated control (FDR<0.05; log_2_FC>1 or <-1).

### Gene annotation and gene ontology enrichment analysis

Differentially expressed genes were annotated using soybean genome gene annotations (Annotation_Gmax_275_Wm82.a2.v1.gene.gff3) from Phytozome (https://phytozome.jgi.doe.gov/pz/portal.html#!bulk?org=Org_Gmax) while soybean chloroplast (https://www.ncbi.nlm.nih.gov/nuccore/91214122) and mitochondrion (https://www.ncbi.nlm.nih.gov/nuccore/476507670) sequences were used from NCBI. The statistically significant DEGs (p-value<0.01; log_2_FC>1 or <-1) of the R v S comparison were used to identify enriched Gene ontology (GO) terms using the Soybase GO term enrichment tool (https://www.soybase.org/goslimgraphic_v2/dashboard.php) (34).

### Metabolites estimation and analysis

Four-week-old soybean plants of susceptible and resistant lines were petiole inoculated as described above. Four stems (3 cm) for each treatment per biological replicate were harvested 48 and 72 hours post inoculation and non-inoculated stems were used as a control. Collected stem samples were immediately frozen in liquid nitrogen and kept at –80°C until used. Gas chromatographic mass spectrometry (GC-MS) analysis was performed by West Coast Metabolomics (Univ. California, Davis). For metabolite estimation, stem tissue from each treatment was finely ground with liquid nitrogen and extracted with 1 ml of chilled 5:2:2 MeOH: CHCl_3_: H2O. The aqueous phase was transferred into a new tube and a 500 μl aliquot was dried under vacuum and the residue was derivatized in a final volume of 100 μl. Detailed methods of metabolite derivatization, separation, and detection are described in Fiehn et al., 2016 (97). Briefly, samples were injected (0.5 μl, split less injection) into a Pegasus IV GC (Leco Corp., St Joseph, MI) equipped with a 30 m x 0.25 mm i.d. fused-silica capillary column bound with 0.25 μm Rtx-5Sil MS stationary phase (Restek Corporation, Bellefonte, PA). The injector temperature was 50°C ramped to 250°C by 12°C s-1. A helium mobile phase was applied at a flow rate of 1 ml min^−1^. Column temperature was 50°C for 1 min, ramped to 330°C by 20°C min-1 and held constant for 5 min. The column effluent was introduced into the ion source of a Pegasus IV TOF MS (Leco Corp., St Joseph, MI) with transfer line temperature set to 230°C and ion source set to 250°C. Ions were generated with a −70 eV and 1800 V ionization energy. Masses (80-500 m/z) were acquired at a rate of 17 spectra s-1. ChromaTOF 2.32 software (Leco Corp) was used for automatic peak detection and deconvolution using a 3 s peak width. Peaks with signal/noise below 5:1 were rejected. Metabolites were quantified by peak height for the quantification ion. Metabolites were annotated with the BinBase 2.0 algorithm (98).

Statistical analyses were conducted in MetaboAnalyst 3.0 (35). Multivariate and univariate statistics were performed on generalized log transformed peak heights. Metabolites with FDR <0.05 were considered differentially regulated. Metabolite pathway analysis was done using the MetaboAnalyst 3.0 Pathway Analysis tool (35).

### Reverse Transcriptase–quantitative PCR (RT-qPCR)

The internodal region of the infected petiole (including symptomatic and non-symptomatic tissue) was used for RNA isolation. Stems (3 cm) were harvested and immediately frozen in liquid nitrogen. RNA was isolated using the above-mentioned protocol and then treated with RNase-free DNase1 (NEB Inc., Ipswich, MA, USA). The RNA was reverse transcribed using the AMV First-Strand cDNA Synthesis Kit (NEB Inc., Ipswitch, MA) and oligo-dT primer according to the manufacturer’s instructions. RT-qPCR was performed using a SensiFAST SYBR No ROX Kit (Bioline USA Inc., Taunton, MA, USA). Each reaction consisted of 5 μL of SensiFAST SYBR No-ROX Mix, 1μL of 1 : 10-fold diluted template cDNAs, and 0.4 μL of 10 μM gene-specific forward and reverse primers in a final volume of 10 μL. Primers were designed using Primer3 software (99,100) for the amplification of gene fragments that were approximately 100 – 200 bp in length and with an annealing temperature of 60°C (Table S7). The primer specificity was checked in silico against the NCBI database through the Primer-BLAST tool (http://www.ncbi.nlm.nih.gov). RT-qPCR was performed on a CFX96 real-time PCR system (Bio-Rad, Hercules, CA). The run conditions were: 2 min of initial denaturation at 95°C; 95°C for 5 s, 58°C for 10 s and 72°C for 20 s (40 cycles). The relative expression of genes was calculated using the 2^-ΔΔCt^ method (101) with the soybean gene GmCon15S (102) as an endogenous control. Three biological replicates were used for each sample.

### Targeted GC-MS analysis of CA, SA, JA, JA-Ile, cis-OPDA, and ABA

Four-week-old soybean plants of susceptible and resistant lines were petiole inoculated with actively growing agar plugs of *S*. *Sclerotiorum*. Four stems (3 cm) for each biological replicate were harvested at 6, 12, 24, 48, and 72 hpi. Uninoculated stems were also collected for the estimation of the basal concentration of the phytohormones. These collected stem samples were immediately frozen with liquid nitrogen and kept at –80°C until used. Stem tissues from each treatment were finely ground with liquid nitrogen and 100 mg of the ground tissue was added to 500 μl of phytohormone extraction buffer (1-propanol/water/HCl [2:1:0.002 vol/vol/v]) and 10 μl of 5 μM solution of deuterated internal standards: d-ABA ([2H6](+)-cis,trans-ABA; [Olchem]), d-IAA ([2H5] indole-3-acetic acid, Olchem), d-JA (2,4,4-d3; acetyl-2,2-d2 JA; CDN Isotopes), and d-SA (d6-SA, Sigma) and analyzed using GC-MS (103–105). The simultaneous detection of several hormones was accomplished using the methods described in Muller et al 2011 with modifications (106). The analysis utilized an Ascentis Express C-18 Column (3 cm × 2.1 mm, 2.7 μm) connected to an API 3200 LC-electrospray ionization-tandem mass spectrometry (MS/ MS) with multiple reaction mentoring (MRM). The injection volume was 2 μl and had a 600 μl/min mobile phase consisting of Solution A (0.05% acetic acid in water) and Solution B (0.05% acetic acid in acetonitrile) with a gradient consisting of (time – %B): 0.3 – 1%, 2 – 45%, 5 – 100%, 8 – 100%, 9 – 1%, 11 – stop. Three biological replicates of each treatment were performed.

### Plate inhibition assay of *S*. *sclerotiorum*

The plate growth inhibition assay of *S*. *sclerotiorum* was done on solid PDA culture plates containing 0, 250, 500, or 1000 μg/mL of ferulic or caffeic acid. Three replicates were used for each treatment. Plates were inoculated with an actively growing plug of *S*. *sclerotiorum* and grown at 25 °C for either 48 hours (ferulic acid) or 7 days (caffeic acid) prior to assessment.

### Compound extraction from soybean stem

Five hundred mg of infected red stem or unaffected green stem was mixed 1:1 w/v in 100% ethanol at 80°C for 1 h. Samples were resuspended with 100 μl of DMSO and used for *S*. *sclerotiorum* inhibition assay, chemical genomics, and cell permeability assays.

### Chemical genomic analysis

Chemical genomic analysis of the red stem extract was performed using the non-essential yeast deletion mutant collection as described previously (53). Briefly, triplicate 200 μL cultures of the pooled deletion collection were exposed to a 1:10 dilution of the red stem extract and allowed to grow for 48 h. Genomic DNA was extracted using the Invitrogen Purelink 96-well genomic extraction kit (Invitrogen, Carlsbad, CA, USA). Mutant specific barcodes were amplified using indexed primers. Samples were sequenced on a HiSeq2500 (1X50bp) (Illumina, San Diego CA, USA) rapid run and reads were processed using BEAN-counter (107) and EdgeR (108).

### Cell permeability assay

To quantify the membrane damage caused by the red stem extract a FungaLight^™^ Cell Viability assay (Invitrogen L34952) and Guava Flow Cytometer (Millipore, USA) was used as described previously (109). Amphotericin B (100 μg/mL) and poacic acid (100 μg/mL) were included as positive controls. Fluconazole (1 mg/mL) was also included as a control, given its ability to inhibit ergosterol biosynthesis without causing rapid cell permeability. We exposed 200 μL log phase populations of yeast cells in YPD media to the control drugs, red stem extract (0.01, 0.005, and 0.0025%), the control green stem extract at (0.01, 0.005, and 0.0025%), and a 1% DMSO control (n=3) for 4 h at 30°C. The cells were then stained with the FungaLight^™^ kit and immediately analyzed by flow cytometry.

### Biomass and ergosterol estimation of *S*. *sclerotiorum*

The biomass of *S*. *sclerotiorum* was measured by growing the fungus in potato dextrose broth (PDB). Freshly grown PDA cultures were scraped and then washed twice with water at 4000 rpm (4°C) before being resuspended in water. Equal amounts of resuspended *S*. *sclerotiorum* were inoculated into 250 mL conical flasks containing 30 mL PDB. For each 30 ml of PDB, 300μl of red stem extract, green stem extract, or DMSO were added and incubated over a period of 48 hours. To estimate the wet weight, the mycelia were filtered on a pre-weighed Miracloth (Darmstadt, Germany). Ergosterol level was estimated as described by Yarden et. al 2014 (57) and Arthington-Skaggs et al 1999 (110). Briefly, mycelia of the *S*. *sclerotiorum* treated with either red stem extract, green stem extract, or DMSO as described above were washed with distilled water. One gram of mycelium for each strain was resuspended in 3 ml of 25% alcoholic potassium hydroxide solution and vortexed vigorously. Mycelia were then incubated at 85°C for 1 h and sterols extracted by adding 4 ml of n-heptane solution (25% sterile distilled water and 75% n-heptane). The heptane layer was transferred to a clean glass tube and spectrophotometric readings taken between 230 and 300 nm. Four biological replicates were used for each treatment. Ergosterol content was calculated as a percentage/g of wet weight using the following equations: % ergosterol + % 24(28)DHE = [(A_281.5_/290) × F], % 24(28)DHE = [(A_230_/518) × F], where F is the factor for dilution in ethanol and 290 and 518 are the E values determined for crystalline ergosterol and 24(28) dehydroergosterol (DHE), respectively.

## Acknowledgments

We thank Grace Brunette and Haley Mesick for their technical assistance. We thank Professor Caitilyn Allen and her group for stimulating discussions. Next-generation sequencing was performed by the Biotechnology Center at the University of Wisconsin-Madison.

## Supplementary Figure Legends

**Supplementary Figure 1. Partial least squares-discriminate analysis (PLS-DA) score plots of metabolic profiles in soybean R and S line**. The first (PC1) and second (PC2) principal components explain 52.7% of the variance. Control samples are R0 and S0. *S*. *sclerotiorum* infected samples at 24, 48 and 96 hpi for the R lines are represented as R24, R48, and R96, respectively. *S*. *sclerotiorum* infected samples at 24, 48 and 96 hpi for the S line are represented as S24, S48, and S96, respectively. Numbers 1, 2, and 3 represents three independent biological replicates.

**Supplementary Figure 2. Effect of Ferulic and Caffeic acids on *S*. *sclerotiorum* growth**. Ferulic acid (A) inhibits the growth of *S*. *sclerotiorum*. Representative photographs were taken 48 hours post inoculation. Caffeic acid (B) effects normal development of *S*. *sclerotiorum*. Representative photographs were taken 5 days post inoculation. DMSO (Dimethyl sulfoxide) is the solvent control.

**Supplementary Figure 3**. (A) Increased accumulation of the jasmonic acid precursor linolenic acid in the R line compared to the S line, (B) Increased accumulation of cyanoalanine (an indicator of ethylene biosynthesis) in the R line compared to the S line. The bars represent the standard deviation (n = 3). ^∗^ Indicates a significantly difference at p-value < 0.05 (t-test).

**Supplementary Figure 4**. (A) Red and green stem extract of a R line plant infected with *S*. *sclerotiorum* and a S line plant mock inoculated. Extraction was performed 10 dpi, (B) Fungal biomass after growth in PDB cultures containing the red stem extract, DMSO, or green stem extract, (C) weight of fungal biomass in PDB cultures containing the red stem extract, green stem extract, or DMSO.

## List of supplementary tables

Supplementary Table 1. Differentially expressed genes (DEGs) in R and S lines following *S*. *sclerotiorum* infection at 24, 48, and 96 hpi compared to control. (A-F) Comparisons between each time-point for both lines and their respective controls.

Supplementary Table 2. Differentially expressed genes in the R line compared to the S line following *S*. *sclerotiorum* infection at 24, 48, and 96 hpi. (A-C) Time-point comparisons between R and S.

Supplementary Table 3. GO enrichment of significant biological processes generated from differentially regulated genes in the R line compared to the S line. (A-C) GO processes identified through the comparison of the R and S lines at 24, 48, and 96 hpi, (D) Summary of significantly regulated genes belonging to each process at each time-point. Processes were included if they had a FDR<0.05 in at-least one timepoint.

Supplementary Table 4. Estimated gas chromatography – mass spectrometry (GC-MS) peak intensity list of all the metabolites.

Supplementary Table 5. Significantly regulated metabolites in the R line compared to the S line following *S*. *sclerotiorum* infection at 24, 48, and 96 hpi. Blue = upregulated. Red = downregulated.

Supplementary Table 6. Metabolic pathways assigned to significantly regulated metabolites from comparison of R and S lines at 48 and 72 hpi. (A) Percentage of all annotated metabolites within each pathway which were found to be significantly regulated in this study. Upregulated = Green. Downregulated = Red, (B) Fold changes of the individual metabolites assigned to each pathway. Values >1 demonstrate upregulation. Value <1 demonstrate downregulation. Values =1 demonstrate no change.

Supplementary Table 7. Primer list for qRT-PCR of phenylpropanoid genes

Supplementary Table 8. Differentially expressed genes encoding putative reactive oxygen species (ROS) scavenging and antioxidant genes in the R line compared to the S line following *S*. *sclerotiorum* infection at 24, 48, and 96 hpi.

Supplementary Table 9. Differentially expressed genes encoding putative jasmonic acid (JA) and ethylene (ET) biosynthetic and response genes in the R line compared to the S line following *S*. *sclerotiorum* infection at 24, 48, and 96 hpi.

## References

1. Boland GJ, Hall R. Index of plant hosts of sclerotinia sclerotiorum. Can J Plant Pathol. 1994;16(2):93–108.

2. Baker DG, Burkey AKO, Burton A, Carter TE, Orf JH, Bernardo R, et al. Soybean Pathology White Paper. Crop Sci [Internet]. 2010;103(2):/PHP-2010-1122-01-RS. Available from: https://www.crops.org/publications/cm/abstracts/6/1/0%5Cnhttp://www.pubmedcentral.nih.gov/articlerender.fcgi?artid=3272368&tool=pmcentrez&rendertype=abstract%5Cnhttp://www.pubmedcentral.nih.gov/articleadaprender.fcgi?artid=2586459&tool=pmcentrez&rendertype=ab

3. Peltier AJ, Bradley CA, Chilvers MI, Malvick DK, Mueller DS, Wise KA, et al. Biology, Yield loss and Control of Sclerotinia Stem Rot of Soybean. J Integr Pest Manag [Internet]. 2012;3(2):1–7. Available from: https://academic.oup.com/jipm/article-lookup/doi/10.1603/IPM11033

4. Cunha WG, Tinoco MLP, Pancoti HL, Ribeiro RE, Aragão FJL. High resistance to Sclerotinia sclerotiorum in transgenic soybean plants transformed to express an oxalate decarboxylase gene. Plant Pathol. 2010;59(4):654–60.

5. Hill J, Nelson E, Tilman D, Polasky S, Tiffany D. Environmental, economic, and energetic costs and benefits of biodiesel and ethanol biofuels. Proc Natl Acad Sci [Internet]. 2006;103(30):11206–10. Available from: http://www.pnas.org/cgi/doi/10.1073/pnas.0604600103

6. Ribeiro MDMM, Ming CC, Lopes TIB, Grimaldi R, Marsaioli AJ, Gonçalves LAG. Synthesis of structured lipids containing behenic acid from fully hydrogenated Crambe abyssinica oil by enzymatic interesterification. J Food Sci Technol. 2017;54(5):1146–57.

7. Hartman GL, West ED, Herman TK. Crops that feed the World 2. Soybean-worldwide production, use, and constraints caused by pathogens and pests. Food Secur. 2011;3(1):5–17.

8. Hoffman DD, Hartman GL, Mueller DS, Leitz RA, Nickell CD, Pedersen WL. Yield and Seed Quality of Soybean Cultivars Infected with Sclerotinia sclerotiorum. Plant Dis [Internet]. 1998;82(7):826–9. Available from: http://apsjournals.apsnet.org/doi/abs/10.1094/PDIS.1998.82.7.826

9. McCaghey M, Willbur J, Ranjan A, Grau CR, Chapman S, Diers B, et al. Development and Evaluation of Glycine max Germplasm Lines with Quantitative Resistance to Sclerotinia sclerotiorum. Front Plant Sci [Internet]. 2017;8(August):1–13. Available from: http://journal.frontiersin.org/article/10.3389/fpls.2017.01495/full

10. Amselem J, Cuomo CA, van Kan JAL, Viaud M, Benito EP, Couloux A, et al. Genomic analysis of the necrotrophic fungal pathogens sclerotinia sclerotiorum and botrytis cinerea. PLoS Genet. 2011;7(8).

11. Kabbage M, Williams B, Dickman MB. Cell Death Control: The Interplay of Apoptosis and Autophagy in the Pathogenicity of Sclerotinia sclerotiorum. PLoS Pathog. 2013;9(4).

12. Liang X, Liberti D, Li M, Kim YT, Hutchens A, Wilson R, et al. Oxaloacetate acetylhydrolase gene mutants of Sclerotinia sclerotiorum do not accumulate oxalic acid, but do produce limited lesions on host plants. Mol Plant Pathol. 2015;16(6):559–71.

13. Xu L, Xiang M, White D, Chen W. pH dependency of sclerotial development and pathogenicity revealed by using genetically defined oxalate-minus mutants of Sclerotinia sclerotiorum. Environ Microbiol. 2015;17(8):2896–909.

14. Williams B, Kabbage M, Kim HJ, Britt R, Dickman MB. Tipping the balance: Sclerotinia sclerotiorum secreted oxalic acid suppresses host defenses by manipulating the host redox environment. PLoS Pathog. 2011;7(6).

15. Kim KS, Min J, Dickman MB. Oxalic Acid Is an Elicitor of Plant Programmed Cell Death during Sclerotinia sclerotiorum Disease Development. 2008;21(5):605–12.

16. Ranjan A, Jayaraman D, Grau C, Hill JH, Whitham SA, Ané JM, et al. The pathogenic development of Sclerotinia sclerotiorum in soybean requires specific host NADPH oxidases. Mol Plant Pathol. 2018;19(3):700–14.

17. Dallal Bashi Z, Hegedus DD, Buchwaldt L, Rimmer SR, Borhan MH. Expression and regulation of Sclerotinia sclerotiorum necrosis and ethylene-inducing peptides (NEPs). Mol Plant Pathol. 2010;11(1):43–53.

18. Nováková M, Sa V, Dobrev PI, Valentová O, Burketová L. Plant Physiology and Biochemistry Plant hormones in defense response of Brassica napus to Sclerotinia sclerotiorum e Reassessing the role of salicylic acid in the interaction with a necrotroph. 2014;80:308–17.

19. Zhu W, Wei W, Fu Y, Cheng J, Xie J, Li G, et al. A Secretory Protein of Necrotrophic Fungus Sclerotinia sclerotiorum That Suppresses Host Resistance. PLoS One. 2013;8(1).

20. Pedras MSC, Ahiahonu PWK. Phytotoxin production and phytoalexin elicitation by the phytopathogenic fungus Sclerotinia sclerotiorum. J Chem Ecol. 2004;30(11):2163–79.

21. Grant D, Nelson RT, Cannon SB, Shoemaker RC. SoyBase, the USDA-ARS soybean genetics and genomics database. Nucleic Acids Res. 2009;38(SUPPL.1):843–6.

22. Kim HS, Diers BW. Inheritance of partial resistance to sclerotinia stem rot in soybean. Crop Sci. 2000;40(1):55–61.

23. Arahana VS, Graef GL, Specht JE, Steadman JR, Eskridge KM. Identification of QTLs for Resistance to in Soybean. Crop Sci. 2001;41(1):180.

24. Huynh TT, Bastien M, Iquira E, Turcotte P, Belzile F. Identification of QTLs associated with partial resistance to white mold in soybean using field-based inoculation. Crop Sci. 2010;50(3):969–79.

25. Moellers TC, Singh A, Zhang J, Brungardt J, Kabbage M, Mueller DS, et al. Main and epistatic loci studies in soybean for Sclerotinia sclerotiorum resistance reveal multiple modes of resistance in multi-environments. Sci Rep. 2017;7(1):1–13.

26. Wei W, Mesquita ACO, Figueiró A de A, Wu X, Manjunatha S, Wickland DP, et al. Genome-wide association mapping of resistance to a Brazilian isolate of Sclerotinia sclerotiorum in soybean genotypes mostly from Brazil. BMC Genomics. 2017;18(1):1–16.

27. Joshi RK, Megha S, Basu U, Rahman MH, Kav NNV. Genome wide identification and functional prediction of long non-coding RNAs responsive to Sclerotinia sclerotiorum infection in Brassica napus. PLoS One. 2016;11(7):1–19.

28. Girard IJ, Tong C, Becker MG, Mao X, Huang J, De Kievit T, et al. RNA sequencing of Brassica napus reveals cellular redox control of Sclerotinia infection. J Exp Bot. 2017;68(18):5079–91.

29. Seifbarghi S, Borhan MH, Wei Y, Coutu C, Robinson SJ, Hegedus DD. Changes in the Sclerotinia sclerotiorum transcriptome during infection of Brassica napus. BMC Genomics. 2017;18(1):1–37.

30. Zhuang X, McPhee KE, Coram TE, Peever TL, Chilvers MI. Rapid transcriptome characterization and parsing of sequences in a non-model host-pathogen interaction; pea-Sclerotinia sclerotiorum. BMC Genomics. 2012;13(1):1–19.

31. Oliveira MB, de Andrade R V., Grossi-de-Sá MF, Petrofeza S. Analysis of genes that are differentially expressed during the Sclerotinia sclerotiorum-Phaseolus vulgaris interaction. Front Microbiol. 2015;6(OCT):1–14.

32. Peltier AJ, Hatfield RD, Grau CR. Soybean stem lignin concentration relates to resistance to Sclerotinia sclerotiorum. Plant Dis. 2009;93:149–54.

33. Goodstein DM, Shu S, Howson R, Neupane R, Hayes RD, Fazo J, et al. Phytozome: A comparative platform for green plant genomics. Nucleic Acids Res. 2012;40(D1):1178–86.

34. Morales AMAP, van de Mortel M, E TJB, a AB, F RTN, E DN, et al. Transcriptome analyses and virus induced gene silencing identify genes in the Rpp4 -mediated Asian soybean rust resistance pathway. Funct Plant Biol. 2013;40:1029–47.

35. Xia J, Sinelnikov I V., Han B, Wishart DS. MetaboAnalyst 3.0-making metabolomics more meaningful. Nucleic Acids Res. 2015;43(W1):W251–7.

36. Wasternack C, Song S. Jasmonates: Biosynthesis, metabolism, and signaling by proteins activating and repressing transcription. J Exp Bot. 2017;68(6):1303–21.

37. Yip WK, Yang SF. Ethylene biosynthesis in relation to cyanide metabolism. Bot Bull Acad Sin [Internet]. 1998;39(1):1–7. Available from: http://ejournal.sinica.edu.tw/bbas/content/1998/1/bot91-01.pdf

38. Naoumkina MA, Zhao Q, Gallego-Giraldo L, Dai X, Zhao PX, Dixon RA. Genome-wide analysis of phenylpropanoid defence pathways. Mol Plant Pathol. 2010;11(6):829–46.

39. Piasecka A, Jedrzejczak-Rey N, Bednarek P. Secondary metabolites in plant innate immunity: Conserved function of divergent chemicals. New Phytol. 2015;206(3):948–64.

40. Widhalm JR, Dudareva N. A familiar ring to it: Biosynthesis of plant benzoic acids. Mol Plant [Internet]. 2015;8(1):83–97. Available from: http://dx.doi.org/10.1016/j.molp.2014.12.001

41. Sharma P, Jha AB, Dubey RS, Pessarakli M. Reactive Oxygen Species, Oxidative Damage, and Antioxidative Defense Mechanism in Plants under Stressful Conditions. J Bot [Internet]. 2012;2012:1–26. Available from: http://www.hindawi.com/journals/jb/2012/217037/

42. Apel K, Hirt H. REACTIVE OXYGEN SPECIES: Metabolism, Oxidative Stress, and Signal Transduction. Annu Rev Plant Biol [Internet]. 2004;55(1):373–99. Available from: http://www.annualreviews.org/doi/10.1146/annurev.arplant.55.031903.141701

43. Hayat S, Hayat Q, Alyemeni MN, Wani AS, Pichtel J, Ahmad A. Role of proline under changing environments: A review. Plant Signal Behav. 2012;7(11).

44. Kim KH, Jia B, Jeon CO. Identification of trans-4-hydroxy-L-proline as a compatible solute and its biosynthesis and molecular characterization in Halobacillus halophilus. Front Microbiol. 2017;8(OCT): 1–11.

45. Smirnoff N, Cumbes QJ. Hydroxyl radical scavening activity of compatible solutes. Phytochimistry. 1989;28(4):1057–60.

46. Matysik J, Alia, Bhalu B, Mohanty P. Molecular mechanisms of quenching of reactive oxygen species by proline under stress in plants. Curr Sci. 2002;82(5):525–32.

47. Bari R, Jones JDG. Role of plant hormones in plant defence responses. Plant Mol Biol. 2009;69(4):473–88.

48. Pieterse CMJ, Van der Does D, Zamioudis C, Leon-Reyes A, Van Wees SCM. Hormonal Modulation of Plant Immunity. Annu Rev Cell Dev Biol [Internet]. 2012;28(1):489–521. Available from: http://www.annualreviews.org/doi/10.1146/annurev-cellbio-092910-154055

49. Robert-Seilaniantz A, Grant M, Jones JDG. Hormone Crosstalk in Plant Disease and Defense: More Than Just JASMONATE-SALICYLATE Antagonism. Annu Rev Phytopathol [Internet]. 2011;49(1):317–43. Available from: http://www.annualreviews.org/doi/10.1146/annurev-phyto-073009-114447

50. Pieterse CMJ, Leon-Reyes A, Van Der Ent S, Van Wees SCM. Networking by small-molecule hormones in plant immunity. Nat Chem Biol. 2009;5(5):308–16.

51. Veronese P, Chen X, Bluhm B, Salmeron J, Dietrich R, Mengiste T. The BOS loci of Arabidopsis are required for resistance to Botrytis cinerea infection. Plant J. 2004;40(4):558–74.

52. Glazebrook J. Contrasting Mechanisms of Defense Against Biotrophic and Necrotrophic Pathogens. Annu Rev Phytopathol [Internet]. 2005;43(1):205–27. Available from: http://www.annualreviews.org/doi/10.1146/annurev.phyto.43.040204.135923

53. Piotrowski JS, Simpkins SW, Li SC, Deshpande R, Ong I, Myers CL, et al. Chemical Biology. 2015;1263:1–19. Available from: http://link.springer.com/10.1007/978-1-4939-2269-7

54. Costanzo M, VanderSluis B, Koch EN, Baryshnikova A, Pons C, Tan G, et al. A global genetic interaction network maps a wiring diagram of cellular function. Science (80-). 2016;353(6306).

55. Pascual F, Soto-Cardalda A, Carman GM. PAH1-encoded phosphatidate phosphatase plays a role in the growth phase- and inositol-mediated regulation of lipid synthesis in Saccharomyces cerevisiae. J Biol Chem. 2013;288(50):35781–92.

56. Parsons AB, Lopez A, Givoni IE, Williams DE, Gray CA, Porter J, et al. Exploring the Mode-of-Action of Bioactive Compounds by Chemical-Genetic Profiling in Yeast. Cell. 2006;126(3):611–25.

57. Yarden O, Veluchamy S, Dickman MB, Kabbage M. Sclerotinia sclerotiorum catalase SCAT1 affects oxidative stress tolerance, regulates ergosterol levels and controls pathogenic development. Physiol Mol Plant Pathol [Internet]. 2014;85:34–41. Available from: http://dx.doi.org/10.1016/j.pmpp.2013.12.001

58. Tripathy BC hara., Oelmüller R. Reactive oxygen species generation and signaling in plants. Plant Signal Behav. 2012;7(12):1621–33.

59. Waszczak C, Carmody M. Reactive Oxygen Species in Plant Signaling. 2009;(February):1–28. Available from: http://link.springer.com/10.1007/978-3-642-00390-5

60. Zurbriggen MD, Carrillo N, Hajirezaei MR. ROS signaling in the hypersensitive response: When, where and what for? Plant Signal Behav. 2010;5(4):393–6.

61. Keinath NF, Kierszniowska S, Lorek J, Bourdais G, Kessler SA, Shimosato-Asano H, et al. PAMP (Pathogen-associated Molecular Pattern)-induced changes in plasma membrane compartmentalization reveal novel components of plant immunity. J Biol Chem. 2010;285(50):39140–9.

62. Gilbert BM, Wolpert TJ. Characterization of the LOV1-mediated, victorin-induced, cell-death response with virus-induced gene silencing. Mol Plant Microbe Interact [Internet]. 2013;26(8):903–17. Available from: http://www.ncbi.nlm.nih.gov/pubmed/23634836

63. Wei L, Jian H, Lu K, Filardo F, Yin N, Liu L, et al. Genome-wide association analysis and differential expression analysis of resistance to Sclerotinia stem rot in Brassica napus. Plant Biotechnol J. 2016;14(6):1368–80.

64. Wen L, Tan TL, Shu J Bin, Chen Y, Liu Y, Yang ZF, et al. Using proteomic analysis to find the proteins involved in resistance against Sclerotinia sclerotiorum in adult Brassica napus. Eur J Plant Pathol. 2013;137(3):505–23.

65. Garg H, Li H, Sivasithamparam K, Barbetti MJ. Differentially Expressed Proteins and Associated Histological and Disease Progression Changes in Cotyledon Tissue of a Resistant and Susceptible Genotype of Brassica napus Infected with Sclerotinia sclerotiorum. PLoS One. 2013;8(6).

66. Mei J, Ding Y, Li Y, Tong C, Du H, Yu Y, et al. Transcriptomic comparison between Brassica oleracea and rice (Oryza sativa) reveals diverse modulations on cell death in response to Sclerotinia sclerotiorum. Sci Rep [Internet]. 2016;6(February):1–11. Available from: http://dx.doi.org/10.1038/srep33706

67. Foyer CH. Redox Homeostasis and Antioxidant Signaling: A Metabolic Interface between Stress Perception and Physiological Responses. Plant Cell Online [Internet]. 2005;17(7):1866–75. Available from: http://www.plantcell.org/cgi/doi/10.1105/tpc.105.033589

68. Foyer CH, Noctor G. Oxidant and antioxidant signaling in plants: a re-evaluation of the concept of oxidative stress in a physiological context. Plant, Cell Environ. 2005;28:1056–1071.

69. Wilson JX, Wilson JX. The physiological role of dehydroascorbic acid. FEBS Lett. 2002;527(1–3):5–9.

70. Agarwal A, Kaul V, Faggian R, Rookes JE, Ludwig-müller J, Cahill DM. Analysis of global host gene expression during the primary phase of the Arabidopsis thaliana – Plasmodiophora brassicae interaction. Funct Plant Biol [Internet]. 2011;38(6):462–78. Available from: http://dx.doi.org/10.1071/FP11026%5Cnhttp://www.publish.csiro.au/?act=view_file&file_id=FP11026.pdf

71. Gravot A, Deleu C, Wagner G, Lariagon C, Lugan R, Todd C, et al. Arginase induction represses gall development during clubroot infection in arabidopsis. Plant Cell Physiol. 2012;53(5):901–11.

72. Yan C, Xie D. Jasmonate in plant defence: Sentinel or double agent? Plant Biotechnol J. 2015;13(9):1233–40.

73. Pauwels L, Goossens A. The JAZ Proteins: A Crucial Interface in the Jasmonate Signaling Cascade. Plant Cell Online [Internet]. 2011;23(9):3089–100. Available from: http://www.plantcell.org/cgi/doi/10.1105/tpc.111.089300

74. Herrera-VÃ¡squez A, Salinas P, Holuigue L. Salicylic acid and reactive oxygen species interplay in the transcriptional control of defense genes expression. Front Plant Sci [Internet]. 2015;6(March):1–9. Available from: http://www.frontiersin.org/Plant-Microbe_Interaction/10.3389/fpls.2015.00171/abstract

75. Gundlach H, Müller MJ, Kutchan TM, Zenk MH. Jasmonic acid is a signal transducer in elicitor-induced plant cell cultures. Proc Natl Acad Sci U S A [Internet]. 1992;89(6):2389–93. Available from: http://www.pubmedcentral.nih.gov/articlerender.fcgi?artid=48663&tool=pmcentrez&rendertype=abstract

76. Mueller MJ, Brodschelm W, Spannagl E, Zenk MH. Signaling in the elicitation process is mediated through the octadecanoid pathway leading to jasmonic acid. Proc Natl Acad Sci U S A. 1993;90(16):7490–4.

77. Mirjalili N, Linden JC. Methyl Jasmonate Induced Production of Taxol in Suspension Cultures of. Artif Cells Blood Substitutes Immobil Biotechnol Cells Blood Substit Immobi. 1996;110–8.

78. Brader G. Jasmonate-Dependent Induction of Indole Glucosinolates in Arabidopsis by Culture Filtrates of the Nonspecific Pathogen Erwinia carotovora. Plant Physiol [Internet]. 2001;126(2):849–60. Available from: http://www.plantphysiol.org/cgi/doi/10.1104/pp.126.2.849

79. Fugate KK, de Oliveira LS, Ferrareze JP, Bolton MD, Deckard EL, Finger FL. Jasmonic acid causes short- and long-term alterations to the transcriptome and the expression of defense genes in sugarbeet roots. Plant Gene [Internet]. 2017;9:50–63. Available from: http://dx.doi.org/10.1016/j.plgene.2016.12.006

80. Grabber JH, Ralph J, Hatfield RD. Model studies of ferulate - Coniferyl alcohol crossproduct formation in primary maize walls: Implications for lignification in grasses. J Agric Food Chem. 2002;50(21):6008–16.

81. Vance C. of Disease Resistance ! 1980;(1).

82. Djamei A, Schipper K, Rabe F, Ghosh A, Vincon V, Kahnt J, et al. Metabolic priming by a secreted fungal effector. Nature. 2011;478(7369):395–8.

83. Tanaka S, Brefort T, Neidig N, Djamei A, Kahnt J, Vermerris W, et al. A secreted Ustilago maydis effector promotes virulence by targeting anthocyanin biosynthesis in maize. Elife. 2014;2014(3):1–27.

84. Zhang Y, Butelli E, De Stefano R, Schoonbeek HJ, Magusin A, Pagliarani C, et al. Anthocyanins double the shelf life of tomatoes by delaying overripening and reducing susceptibility to gray mold. Curr Biol [Internet]. 2013;23(12):1094–100. Available from: http://dx.doi.org/10.1016/jxub.2013.04.072

85. Lorenc-Kukuła K, Jafra S, Oszmiański J, Szopa J. Ectopic expression of anthocyanin 5-O-glucosyltransferase in potato tuber causes increased resistance to bacteria. J Agric Food Chem. 2005;53(2):272–81.

86. Kumar N, Pruthi V. Potential applications of ferulic acid from natural sources. Biotechnol Reports [Internet]. 2014;4(1):86–93. Available from: http://dx.doi.org/10.1016/j.btre.2014.09.002

87. Matejczyk M, Świsłocka R, Golonko A, Lewandowski W, Hawrylik E. Cytotoxic, genotoxic and antimicrobial activity of caffeic and rosmarinic acids and their lithium, sodium and potassium salts as potential anticancer compounds. Adv Med Sci. 2018;63(1):14–21.

88. Papadopoulou K, Melton RE, Leggett M, Daniels MJ, Osbourn AE. Compromised disease resistance in saponin-deficient plants. Proc Natl Acad Sci [Internet]. 1999;96(22):12923–8. Available from: http://www.pnas.org/cgi/doi/10.1073/pnas.96.22.12923

89. Bednarek, Pawel; Osbourn A. Plant-microbe interactions: chemical diversity in plant defense. Science [Internet]. 2009;324(5928):746–8. Available from: http://www.ncbi.nlm.nih.gov/pubmed/19423814

90. Simons V, Morrissey JP, Latijnhouwers M, Csukai M, Cleaver A, Yarrow C, et al. Dual effects of plant steroidal alkaloids on Saccharomyces cerevisiae. Antimicrob Agents Chemother. 2006;50(8):2732–40.

91. Rautenbach M, Troskie AM, Vosloo JA. Antifungal peptides: To be or not to be membrane active. Biochimie [Internet]. 2016;130:132–45. Available from: http://dx.doi.org/10.1016/j.biochi.2016.05.013

92. Thevissen K, François IEJA, Takemoto JY, Ferket KKA, Meert EMK, Cammue BPA. DmAMP1, an antifungal plant defensin from dahlia (Dahlia merckii), interacts with sphingolipids from Saccharomyces cerevisiae. FEMS Microbiol Lett. 2003;226(1):169–73.

93. Schmutz J, Cannon SB, Schlueter J, Ma J, Mitros T, Nelson W, et al. Genome sequence of the palaeopolyploid soybean. Nature [Internet]. 2010;463(7278):178–83. Available from: http://dx.doi.org/10.1038/nature08670

94. Liao Y, Smyth GK, Shi W. The Subread aligner: Fast, accurate and scalable read mapping by seed-and-vote. Nucleic Acids Res. 2013;41(10).

95. Law CW, Chen Y, Shi W, Smyth GK. Voom: Precision weights unlock linear model analysis tools for RNA-seq read counts. Genome Biol. 2014;15(2):1–17.

96. Ritchie ME, Phipson B, Wu D, Hu Y, Law CW, Shi W, et al. Limma powers differential expression analyses for RNA-sequencing and microarray studies. Nucleic Acids Res. 2015;43(7):e47.

97. Fiehn O. Metabolomics by gas chromatography-mass spectrometry: Combined targeted and untargeted profiling. Vol. 2016, Current Protocols in Molecular Biology. 2016. 1–32 p.

98. Skogerson K, Wohlgemuth G, Barupal DK, Fiehn O. The volatile compound BinBase mass spectral database. BMC Bioinformatics. 2011;12:1–15.

99. Untergasser A, Cutcutache I, Koressaar T, Ye J, Faircloth BC, Remm M, et al. Primer3-new capabilities and interfaces. Nucleic Acids Res. 2012;40(15):1–12.

100. Koressaar T, Remm M. Enhancements and modifications of primer design program Primer3. Bioinformatics. 2007;23(10):1289–91.

101. Livak KJ, Schmittgen TD. Analysis of relative gene expression data using real-time quantitative PCR and the 2-ΔΔCT method. Methods. 2001;25(4):402–8.

102. Libault M, Thibivilliers S, Bilgin DD, Radwan O, Benitez M, Clough SJ, et al. Identification of Four Soybean Reference Genes for Gene Expression Normalization. Plant Genome J [Internet]. 2008;1(1):44. Available from: https://www.crops.org/publications/tpg/abstracts/171/44

103. Schmelz EA, Engelberth J, Alborn HT, O’Donnell P, Sammons M, Toshima H, et al. Simultaneous analysis of phytohormones, phytotoxins, and volatile organic compounds in plants. Proc Natl Acad Sci [Internet]. 2003;100(18):10552–7. Available from: http://www.pnas.org/cgi/doi/10.1073/pnas.1633615100

104. Schmelz EA, Kaplan F, Huffaker A, Dafoe NJ, Vaughan MM, Ni X, et al. Identity, regulation, and activity of inducible diterpenoid phytoalexins in maize. Proc Natl Acad Sci [Internet]. 2011; 108(13):5455–60. Available from: http://www.pnas.org/cgi/doi/10.1073/pnas.1014714108

105. Christensen SA, Nemchenko A, Park Y-S, Borrego E, Huang P-C, Schmelz EA, et al. The Novel Monocot-Specific 9-Lipoxygenase ZmLOX12 Is Required to Mount an Effective Jasmonate-Mediated Defense Against *Fusarium verticillioides* in Maize. Mol Plant-Microbe Interact [Internet]. 2014;27(11):1263–76. Available from: http://apsjournals.apsnet.org/doi/10.1094/MPMI-06-13-0184-R

106. Müller M, Munné-Bosch S. Rapid and sensitive hormonal profiling of complex plant samples by liquid chromatography coupled to electrospray ionization tandem mass spectrometry. Plant Methods. 2011;7(1):1–11.

107. Piotrowski JS, Li SC, Deshpande R, Simpkins SW, Nelson J, Yashiroda Y, et al. Functional annotation of chemical libraries across diverse biological processes. Nat Chem Biol. 2017;13(9):982–93.

108. Robinson MD, McCarthy DJ, Smyth GK. edgeR: A Bioconductor package for differential expression analysis of digital gene expression data. Bioinformatics. 2009;26(1):139–40.

109. Wyche TP, Piotrowski JS, Hou Y, Braun D, Deshpande R, McIlwain S, et al. Forazoline A: Marine-Derived Polyketide with Antifungal in Vivo Efficacy. Angew Chemie - Int Ed. 2014;53(43):11583–6.

110. Arthington-Skaggs BA, Jradi H, Desai T, Morrison CJ. Quantitation of ergosterol content: Novel method for determination of fluconazole susceptibility of Candida albicans. J Clin Microbiol. 1999;37(10):3332–7.

111. Baxter H, Stewart CN. Effects of altered lignin biosynthesis on phenylpropanoid metabolism and plant stress. Biofuels. 2013;4(6):635–50.

112. Ferrer J-L, Austin MB, Stewart C, Noel JP, Noel JP. Structure and function of enzymes involved in the biosynthesis of phenylpropanoids. Plant Physiol Biochem PPB [Internet]. 2008;46(3):356–70. Available from: http://www.ncbi.nlm.nih.gov/pubmed/18272377%5Cnhttp://www.pubmedcentral.nih.gov/articlerender.fcgi?artid=PMC2860624

